# A user-friendly single-nucleus RNA sequencing pipeline to identify alterations in gene expression in small brain circuits

**DOI:** 10.64898/2026.07.28.741316

**Authors:** GS Stephens, D Alcantara-Gonzalez, HE Scharfman

## Abstract

Single-cell or single-nucleus RNA sequencing are common methods to investigate gene expression. However, to clarify the genes in specific types of cells in a small circuit there are limitations to current approaches. Here we present modifications to standard protocols to overcome the limitations and do so in a manner that will be accessible to novices. Then the modified methods are applied to a question about a small area of the brain, the dentate gyrus (DG) of the mouse, where information about cell types was of interest. The question arose from data acquired in a mouse model of Alzheimer’s disease where early hyperactivity of the principal cells, granule cells (GCs), was identified that was difficult to explain by existing data. Therefore, we investigated altered gene expression in GCs, and other DG cell types that influence GCs, to identify putative mechanisms. Validations of the modified methods are addressed, comparisons are made to other methods, and comparisons of mouse and human data are presented.

## INTRODUCTION

Single-cell and single-nucleus RNA sequencing (scRNA- and snRNA-seq) have revolutionized the study of gene expression in the brain by providing higher resolution and more comprehensive analysis of genes than prior methods. Moreover, these methods are unique in their ability to determine how many RNA transcripts are expressed for each gene within an individual cell or nucleus. The high throughput of these methods can quantify RNA in thousands of cells at once, providing a cell type-specific quantitative atlas of gene expression (Grindberg *et al*., 2013; Cembrowski *et al*., 2016; Walter *et al*., 2023). This atlas therefore provides information on all transcripts expressed in cells in a tissue, often called the transcriptome. Previous transcriptomic methods such as bulk RNA sequencing lacked resolution of single cells and cell type specificity (Grindberg *et al*., 2013). Laser-capture microdissection-sequencing can enhance cell type specificity of bulk sequencing but is often low throughput (Lovatt *et al*., 2015). Spatial transcriptomics allows measurement of gene expression with spatial registration of signals. However, one difficulty is that the raw measurements correspond to locations on a tissue section rather than specific cell types (Satija *et al*., 2015; *preprints:* Salas *et al*., 2025; Hallinan *et al*., 2026). Subsequent computational modeling is required to associate the measurements with individual cell boundaries, which can have limitations (Park *et al*., 2023; Wang *et al*., 2025b; *preprint:* Ludington *et al*., 2026). In contrast, scRNA- or snRNA-seq allows comprehensive analyses of the entire transcriptome in individual cells that are easily differentiated by the use of barcode tags during sequencing (Clark *et al*., 2023; Park *et al*., 2023).

An important question that can be addressed using scRNA- or snRNA-seq emerged from our previous studies of Alzheimer’s disease (AD) using mouse models that simulate features of AD. We found that early in the life of the mice, long before the hallmark pathology and cognitive deficits of AD arose, the brain exhibited hyperactivity. In addition, specific cell types in the dentate gyrus (DG) showed aberrant excitability (Chin and Scharfman, 2013; Duffy *et al*., 2015; Kam *et al*., 2016; Alcantara-Gonzalez *et al*., 2021; Lisgaras and Scharfman, 2023a, b; Chartampila *et al*., 2024; Alcantara-Gonzalez *et al*., 2025). We decided to use scRNA- or snRNA-seq to obtain a comprehensive understanding of potential gene changes in the DG that could cause such early aberrant neuronal activity.

As background, the common perspective of AD mechanisms is that the accumulation of pathological amyloid β (Aβ) and neurofibrillary tau are responsible for loss of memory and behavioral disturbances (Welikovitch *et al*., 2020; Leng *et al*., 2021; Mathys *et al*., 2023; Abdulkhaliq *et al*., 2026). Notably, there are additional contributing factors such as inflammation (Roy *et al*., 2020; Zhou *et al*., 2020; Weaver, 2023; Al-kuraishy *et al*., 2025; Abdulkhaliq *et al*., 2026; Bieri *et al*., 2026; Darvas *et al*., 2026), metabolism (Lyra *et al*., 2021; Naia *et al*., 2023; Hruby and Higuchi-Sanabria, 2025; Hawkinson *et al*., 2026), and abnormalities during sleep (Ju *et al*., 2013; Xie *et al*., 2013; Kam *et al*., 2016; Brown *et al*., 2018; Shokri-Kojori *et al*., 2018; Wang and Holtzman, 2020; Jagirdar *et al*., 2021; Lisgaras and Scharfman, 2023b; Szabo *et al*., 2023; Zavecz *et al*., 2023; Nguyen Ho *et al*., 2024).

New perspectives suggest that hyperactivity of neurons may play a role in AD. This idea began after it was noted that many patients developed abnormal EEG activity, often very early in the disease (Hauser *et al*., 1986; Volicer *et al*., 1995; Amatniek *et al*., 2006; Scarmeas *et al*., 2009). Although initially considered not to be relevant to the majority of patients or contribute to underlying mechanisms, evidence has now accumulated that suggests that hyperactivity is more common and plays a role in AD pathophysiology and memory deficits. Manifestations of hyperactivity have been detected using scalp and intracranial EEG, as well as MEG and fMRI (Bakker *et al*., 2012; Vossel *et al*., 2013; Bakker *et al*., 2015; Vossel *et al*., 2016; Lam *et al*., 2017; Vossel *et al*., 2017; Lam *et al*., 2020; Vöglein *et al*., 2020; Canet *et al*., 2022; Kamondi *et al*., 2024; Nous *et al*., 2024; Devulder *et al*., 2025; Lisgaras *et al*., 2025; Stewart and Johnson, 2025; Devinsky *et al*., 2026). Hyperactivity occurs in both familial and sporadic AD and can be severe, similar to epileptic activity observed in epilepsy (Amatniek *et al*., 2006). Moreover, diverse methods show hyperactivity in many mouse models of AD: EEG, expression of immediate early genes, tests that reveal low thresholds for convulsive seizures, and/or calcium imaging (Palop *et al*., 2007; Sanchez *et al*., 2012; Chin and Scharfman, 2013; Corbett *et al*., 2013; Yamamoto *et al*., 2015; Kam *et al*., 2016; Corbett *et al*., 2017; You *et al*., 2017; Fu *et al*., 2019; Johnson *et al*., 2020; Jagirdar *et al*., 2021; Lisgaras and Scharfman, 2023a, b; Szabo *et al*., 2023; Knox *et al*., 2025; Hegnet *et al*., 2026). Strikingly, treatment with antiseizure medications such as levetiracetam can reduce hyperactivity and improve memory not only in mouse models of AD (Sanchez *et al*., 2012; Corbett *et al*., 2017; Fu *et al*., 2019), but also in AD patients (Cumbo and Ligori, 2010; Bakker *et al*., 2012; Bakker *et al*., 2015; Vossel *et al*., 2021; Hautecloque-Raysz *et al*., 2023; Devinsky *et al*., 2026).

In prior studies, our group and others have shown that brief generalized epileptiform spikes, interictal spikes (IIS), appear at just 4-6 weeks of age (Bezzina *et al*., 2015; Kam *et al*., 2016; Lisgaras and Scharfman, 2023a, b). This is a potentially important observation because it is an age long before memory impairment, or deposition of Aβ as plaques (Hsiao *et al*., 1996). Therefore, IIS may contribute to AD progression.

The initiation site of IIS appears to be the DG based on prior studies (Kam *et al*., 2016; Lisgaras and Scharfman, 2023b). Further studies that used patch clamp electrophysiology at 1 month of age (Alcantara-Gonzalez *et al*., 2021; Alcantara-Gonzalez *et al*., 2025) showed that the principal DG cell type, the granule cell (GC), exhibits increased excitability, as does another DG cell type called the mossy cell (MC). These findings led to the hypothesis that identifying alterations in gene expression of cells within the DG circuit, especially GCs and MCs, could lead to an understanding of potential causes of IIS. Ultimately this information could lead to a strategy to stop IIS.

To investigate differential gene expression in transgenic and wild type mice, we selected the Tg2576 mouse model, the mouse model we had used for patch clamp electrophysiology. The Tg2576 mouse overexpresses a mutated form of amyloid precursor protein (APPSwe) found in a Swedish family with AD (Hsiao *et al*., 1996). As a result, Aβ production is increased and accumulates with age, although at young ages it is primarily oligomeric (soluble, intracellular) rather than plaque (Duffy *et al*., 2015; Criscuolo *et al*., 2024; Alcantara-Gonzalez *et al*., 2025). By 3-4 months of age, memory impairments are evident (Duffy *et al*., 2015) and Aβ plaques are detectable after 6 months of age (Hsiao *et al*., 1996; Kawarabayashi *et al*., 2001; Criscuolo *et al*., 2024).

Using these mice we implemented snRNA-seq at 1 month of age. Data analyses emphasized expression of genes that encode proteins involved in excitability, such as ion channels & transporters, neurotransmitter receptors, and other excitability-related proteins. The choice to use snRNA-seq over scRNA-seq was made because snRNA-seq can be performed on frozen tissue (Krishnaswami *et al*., 2016). Freezing tissue after harvesting introduces a natural stopping point in the protocol. Without this stopping point, fresh cells or nuclei must be processed without delay from euthanasia onward until library preparation, a process that can take over 12-18 hours (Grindberg *et al*., 2013; Bakken *et al*., 2018; Clark *et al*., 2023; Kersey *et al*., 2026). In addition, performing snRNA-seq on frozen tissue allows DG samples to be collected over time in batches and processing to be flexible. Once ready, batches of frozen DG samples can be processed together to mitigate differences between batches (Bakken *et al*., 2018; Clark *et al*., 2023).

As we began to implement snRNA-seq, we encountered several difficulties, including some that would be severe for those without experience, specialized equipment, and/or local sequencing facilities. Although companies exist to process frozen samples, the financial resources that are required can be substantial. Therefore, we developed a processing and analysis pipeline that would be helpful even for inexperienced users (Figures 1A-B and S1).

**Figure 1.**
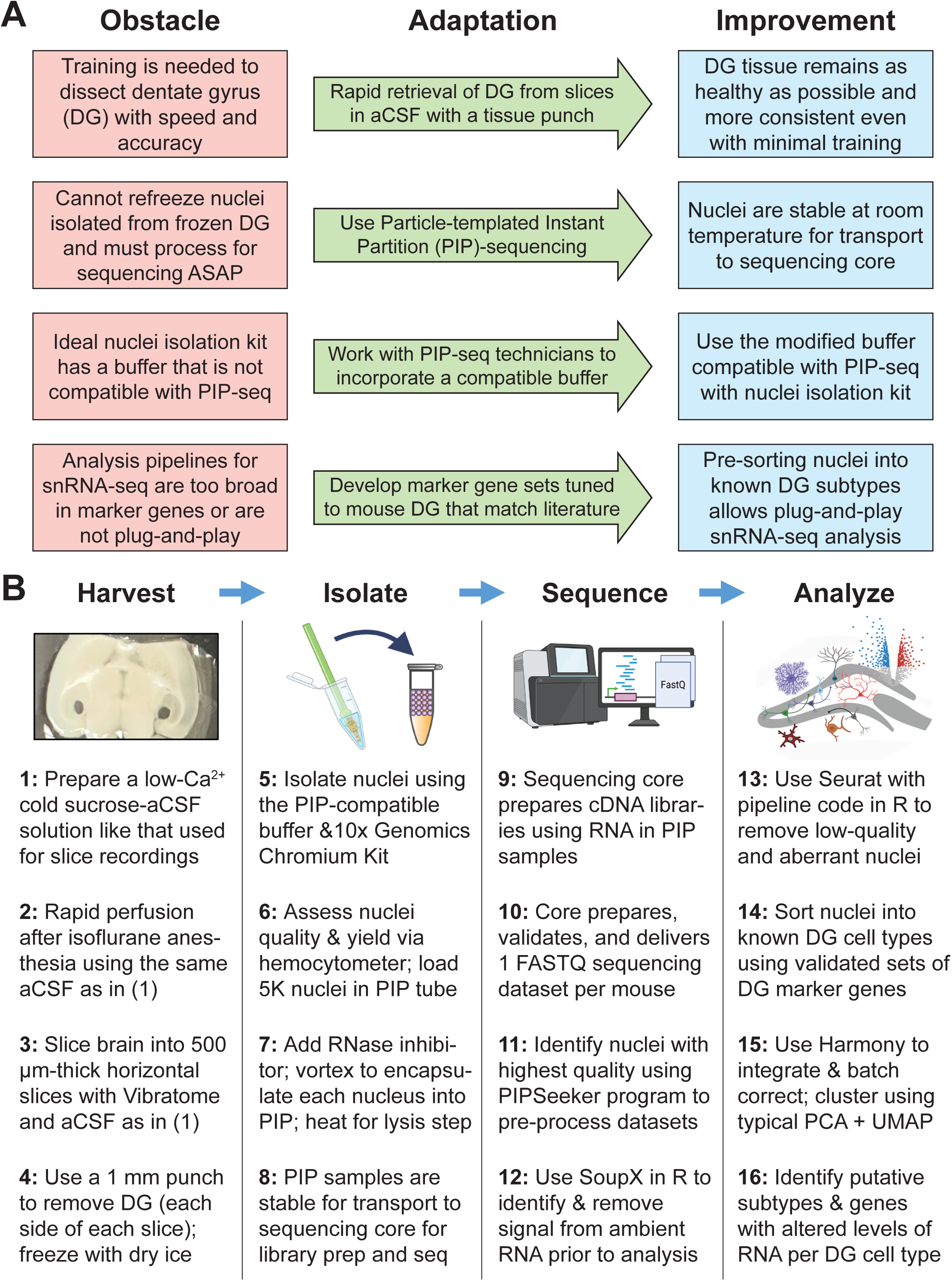
Our user-friendly pipeline circumvents obstacles that can make single-nucleus RNA-sequencing difficult. **(A).** Obstacles for novices (*red*) to perform snRNA-seq can be overcome by modifying existing pipelines (*green*) to enhance efficiency of tissue harvest, stability of samples, compatibility of pipeline reagents, and rigor of analysis (*blue*). **(B).** Our pipeline is composed of 4 primary stages (Figure S1, Methods, and deposited code: https://osf.io/qgty7). The first image in (B) was taken after harvesting DG from a brain slice. An annotated, enlarged version of this image is provided in Figure S2. Other graphics in (B) were created with Biorender.com. Abbreviations: DG, dentate gyrus; aCSF, artificial cerebrospinal fluid; PIP-seq, particle-templated instant partition sequencing; FASTQ, FAST-Quality file format; PCA, principal component analysis; and UMAP, uniform manifold approximation and projection.

In this study, we validate that our snRNA-seq pipeline performs with similar or improved results vs standard methods. Moreover, our pipeline can indeed identify multiple alterations in gene expression in the major cell types in the DG of Tg2576 mice. Furthermore, we show support for our hypothesis that there are alterations in the expression of many genes in multiple DG cell types that may explain how hyperactivity emerges early in life. Also, we compared the single-nucleus gene alterations in our dataset with bulk proteomic alterations in human AD or epilepsy. We confirmed that important gene alterations in multiple DG cell types in young Tg2675 mice can also be detected at the protein level in human AD and epilepsy.

## RESULTS

### I. Our pipeline and the obstacles it overcomes

#### A. Overview

An overview of the pipeline is provided in Figure 1 and details are provided in Figure S1 and Methods. Our pipeline is comparable to existing snRNA-seq pipelines in that it follows similar general phases to generate sequencing data from a tissue (Figure 1B). First is the harvesting phase, requiring the user to remove and freeze fresh tissue. The next step involves the isolation of nuclei and transport to the location where RNA processing and sequencing are performed. Because common methods to isolate nuclei can lead to degradation during transport to a core facility, the core facility is usually at the same or a nearby institution. Next the core facility processes the sample, which involves extracting RNA and then preparing, validating, and sequencing the associated cDNA libraries. The facility sends the dataset to the user, who perform their own validations and removes nuclei that do not meet standards for quality. The user then analyzes their data to identify cell types, differential gene expression, and can establish epigenetic regulatory programs and functional alterations related to their interests. An example of this workflow can be found in Nelson *et al*. (2023) and is discussed in the Results in Section IIC. Our pipeline includes the same steps but makes modifications to the standard methods to overcome several obstacles (below) and make snRNA-seq accessible to novices.

#### B. The obstacles and methods to circumvent them

##### 1. Extracting fresh DG with precision, speed and consistency

One obstacle is that current protocols to isolate the DG from the intact brain have intrinsic limitations. One solution used by others is to implement flow cytometry to concentrate targeted cell types, although this prevents the unbiased capture of all cell populations that are present (Cembrowski *et al*., 2016). Another approach is to pull the DG away from the hippocampus using microdissection tools, but this is associated with damage to the DG. Therefore, it is common for mouse studies to remove the entire hippocampus and not isolate the DG (Nelson *et al*., 2023). However, a consequence of this approach is that the capture of DG cells during nuclei isolation may be diluted by the presence of non-DG cells from the rest of the hippocampus.

Our approach is to initially cool the brain by perfusion with a cold sucrose-based buffer. The buffer is similar to cerebrospinal fluid (CSF) so it is often called artificial CSF (aCSF) but NaCl is replaced by sucrose and concentrations of Ca^2+^ and Mg^2+^ are altered to slow metabolic processes (for details, see Methods). Cold temperature makes the brain become relatively hard and easier to dissect without damage. Next, the brain is rapidly removed and thick brain slices are sectioned with both hemispheres intact using a Vibratome. The brain remains immersed in cold sucrose-based aCSF. Sectioning uses the horizontal plane because the DG is more of an oval in that orientation which makes its extraction with a tissue punch easier. While sections are immersed in cold sucrose-aCSF in a petri dish, a low power microscope is used with the tissue punch to cut out the DG. The punch is a 1 mm-diameter tool used for biopsies (Figures 1B and S1, *blue*). The advantages of this method are the tools are readily available and easily used. In addition, brain slices from control and experimental groups can be cut and maintained under similar conditions. Moreover, the time needed to isolate the DG is short. Further, the procedure maintains tissue integrity better than pulling the DG from the hippocampus. However, there are disadvantages, such as the need for some training and that use of the horizontal plane makes it more difficult to sample the most septal (rostral) part of the DG (discussed further in Methods). Enlarged annotated photographs of the dissected DG area are provided in Figure S2.

##### 2. Sample processing by a sequencing facility should be performed rapidly after nuclei isolation

Once nuclei are isolated from the DG samples (Figures 1B and S1, *yellow*), the samples will degrade over time. The samples cannot be refrozen because cDNA sequencing and library preparation will be reduced in efficacy and quality. Ideally, RNA extraction and library preparation should be performed within a few hours of nuclei isolation. Thus, those steps are often performed by a sequencing service or core facility near the laboratory. This can be a roadblock for laboratories at institutions without a nearby sequencing core facility. What is desirable is a way to maintain stability of samples during transit. Therefore, we used a recent snRNA-seq method that can prevent RNA in nuclei from degrading at ambient temperatures for up to 72 hours. This method is called particle-templated instant partition sequencing (PIP-seq; Fluent Biosciences, now Illumina), which involves labeling single nuclei with unique barcode tags and encasing them in a substance that separates each nucleus from another (Clark *et al*., 2023). This approach is similar to popular droplet-based methods (e.g. 10x Genomics). However, PIP-seq’s partitioning chemistry does not require a specialized piece of equipment (microfluidics device) to capture and label single nuclei. Instead, samples are vortexed to encase single nuclei, barcodes, and RNAse inhibitors into oil-based “PIPs”. This step is essentially an emulsification of nuclei, barcodes and inhibitors. The PIPs are then incubated using dry heat to release RNA from nuclei and stabilize the RNA inside the PIPs. Afterwards, an extended time (72 hours) to transport to a core facility can be allowed without sample degradation (Clark *et al*., 2023).

##### 3. Nuclei isolation from DG for PIP-seq requires specialized equipment and expertise

The step in which nuclei are isolated from frozen DG cells can be difficult for beginners because it requires specialized equipment and experience. Nuclei isolation usually occurs via (1) magnetic separation, in which nuclei are tagged with beads and isolated via magnetic fields, or (2) density gradient centrifugation, in which nuclei are separated from debris using an ultracentrifuge to filter them through a dense solution (often high in sucrose). This procedure isolates the nuclei into a layer below cell debris because nuclei are denser and travel further than debris. Nuclei isolation can also be performed by lysing cells in a tissue homogenizer and using plastic strainers to isolate nuclei. This approach is easy to use and inexpensive. However, it can result in lower nuclei yields than other methods (Fatma and Genevieve, 2018; Ayhan *et al*., 2021; Kersey *et al*., 2026). Therefore, it is not as suitable for recovery of rare cell types or cells from lower tissue volumes (e.g. MCs from DG punches). For all these reasons, we decided to use a kit that is very commonly cited (Chromium Nuclei Isolation kit, 10x Genomics). It isolates nuclei from homogenates of lysed cells using a spin-column tube (Figures 1B and S1, *yellow*). Although there is some experience required, it is a relatively simple procedure compared to the procedures described above. Nuclei are freed from lysed cells and microcentrifuged at 4°C to trap them in columns, where they are washed and then eluted as purified suspensions. However, it uses a suspension buffer incompatible with PIP-seq. To circumvent this issue, we worked with Fluent to modify the nuclei isolation kit to make it compatible (see Methods). Afterwards, standard methods were used for validating quality of data, filtering nuclei, and normalization (Figures 1B and S1; *green*).

##### 4. Existing snRNA-seq pipelines do not readily identify DG-specific cells

To identify cell type-specific alterations in snRNA-seq data, one must first address the high dimensionality of the data, meaning the numerous features or variables in the data. A typical approach is to reduce the dimensionality (dimensional reduction) using Principal Component Analysis (PCA) followed by Uniform Manifold Approximation and Projection (UMAP). For those unfamiliar with this approach, the following briefly explains it (for more detail, see Becht *et al*., 2019; Kiselev *et al*., 2019; Chari and Pachter, 2023; Marx, 2024). Prior to performing PCA, the high number of variables (aka dimensions) in snRNA-seq data correspond to the expression levels of each gene in each nucleus. The number of variables is reduced by condensing them into a smaller set of variables (PCs) that correspond to the most robust sources of variability in the gene expression dataset. An analogous transformation is then performed upon the PCs to reduce the number of variables to 2. This is referred to as “UMAP clustering” of data features with similar variability, where the data features are the clusters. These variables are plotted in 2D coordinates reflecting complex features of the dataset, usually named UMAP 1 and UMAP 2. This plot will show the separations of groups of nuclei with similar variability (again, these are the clusters). Thus, nuclei in the same cluster share similar aspects of variability that distinguish them from other nuclei in other clusters. When used to analyze a high number of cells of distinct types, the clusters should separate into clusters that correspond to different cell types (Chari and Pachter, 2023; Marx, 2024).

However, existing snRNA-seq analysis methods are based on the assumption that a specific cell type can be reproducibly separated from other types into a cluster that has distinct patterns of variability in its gene expression. This assumption is often violated when separating out cell types with low abundance or when clustering with smaller numbers of nuclei (Kiselev *et al*., 2019). This problem can lead to clusters containing nuclei from mixed or unknown cell types (Kiselev *et al*., 2019; Yao *et al*., 2024). Unfortunately, this problem might only be identified after clusters are generated because that is when cell type identification is performed. Cell type identification, often called annotation, is the process that is performed to determine which clusters consist primarily of nuclei that express marker genes that uniquely correspond to specific cell types (Fischer and Gillis, 2021).

There are already standard user-friendly methods that perform cell type annotation after clustering, including online analysis platforms such as PanglaoDB, MapMyCells or scType (Franzén *et al*., 2019; Ianevski *et al*., 2022; *preprint:* Daniel *et al*., 2026). However, a key obstacle we encountered with existing user-friendly annotation tools is that it can be difficult to detect less common cell types, particularly those specific to the DG. As our hypothesis required analyses of uncommon DG cell types, we developed a new snRNA-seq analysis that is tailored to detect these DG cell types. Notably, our approach separates nuclei into cell types *prior* to performing PCA and UMAP clustering, although there are limitations (see Discussion).

Our approach implements a “Suite of Cell Type Sorters” (SOCTS; Figure 2A). To develop SOCTS for the mouse DG we drew upon the extensive mouse DG literature to identify multiple markers of all cell types and then determined genes specific to each cell type. Each set of marker genes was constructed from numerous overlapping sources that have confirmed DG cell type specificity via multiple modalities (such as sn/scRNA-seq, spatial transcriptomics, in situ hybridization (ISH), or immunohistochemistry) including the Allen Brain Institute’s adult mouse ISH database (see Figure 2B and supplemental references in Table S1). For example, MCs were defined as cells in the DG that express *Col19a1, Nmb,* and/or *Calb2*, but do not express *Prox1* or *Vip*. It has been shown that MCs can express *Cck* and/or *Gad1*, genes that are often used to label GABAergic neurons (interneurons). Therefore, we did not use *Cck* or *Gad1* to distinguish MCs from interneurons (see Table S1). Our SOCTS approach was unique because prior snRNA-seq studies of DG often depict UMAP plots that do not have an annotation for MCs specifically (Nelson *et al*., 2023; da Rocha *et al*., 2025). Once nuclei are sorted using SOCTS, the typical downstream analyses can be performed for each cell type, such as detection of differential gene expression between experimental conditions. Notably, after sorting via SOCTS, UMAP clustering can still be performed if desired to identify and analyze putative DG cell subtypes (e.g. dorsal vs ventral, normal vs abnormal; Figure 1B and S1, *tan/black*).

**Figure 2.**
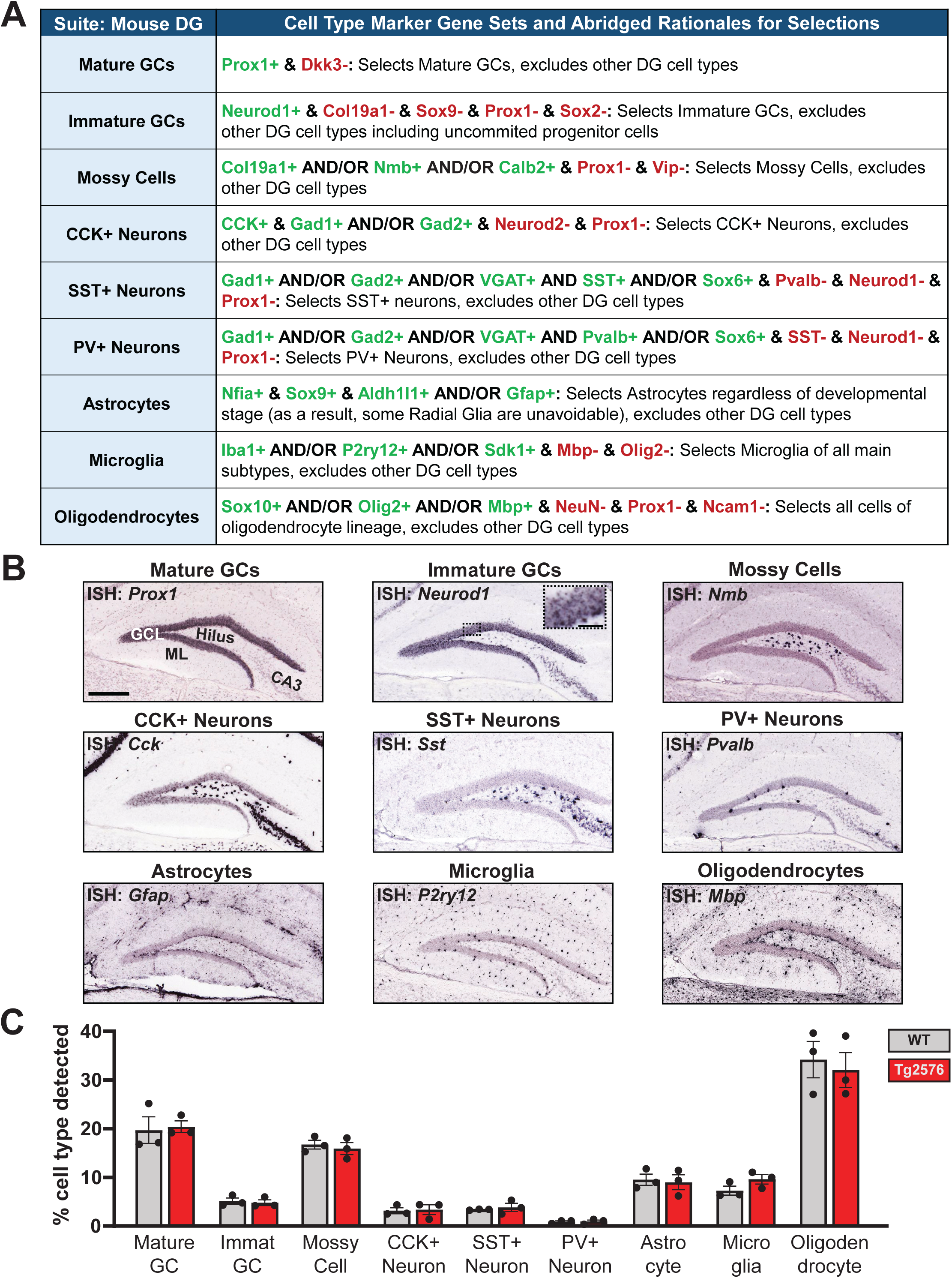
Suite of Cell Type Sorters (SOCTS) uses marker genes to differentiate cell types in the DG prior to clustering or further analysis. **(A).** SOCTS derives its marker gene sets from multiple methods and many publications (listed for each cell type in Table S1). Although other cell types are present in the DG, and our cell types may have subtypes, we chose these 9 major cell types because they are well-characterized, critical to the DG circuit, and are present in sufficient quantities to detect in a single mouse (see C). Positive marker genes that are expressed by a cell type are shown in *green*, with markers that are not expressed by a cell type shown in *red*. **(B).** All sets of marker genes undergo final verification using the Allen Mouse Brain in situ hybridization (ISH) Atlas (Lein *et al*., 2007). ISH data and corresponding hyperlinks are listed in Table S1. **(C).** A two-way ANOVA was performed to identify significant differences in Tg2576 vs wildtype (WT) mice in the proportions of nuclei found for each cell type. Genotype and Cell Type were used as factors. A significant effect of Cell Type was detected (F(8, 36) = 83.73, p < 0.0001), but no effects were found for Genotype (F(1, 36) = 0.000, p > 0.9999) or the interaction (F(8, 36) = 0.2874, p = 0.9658). A Bonferroni post-hoc test corrected for multiple comparisons detected no significant differences in the mean proportions of nuclei detected in DG snRNA-seq samples from Tg2576 vs WT mice (n = 3/group; each p-adj > 0.9999). Raw numbers of nuclei (reported in Figure S3) for each cell type were normalized as percentages to perform the statistics in C because the raw values did not show normality (see Figure S3 legend). Dots = individual mice. Abbreviations: GCL, granule cell layer; ML, molecular layer; GC, granule cell; CCK, cholecystokinin; SST, somatostatin; and PV, parvalbumin. Scale bar = 300 µm. Inset scale bar = 50 µm.

### II. Use of the pipeline to define alterations in specific DG cell types in the Tg2576 model of AD

#### A. Sorting nuclei into DG cell types using SOCTS

Using our pipeline and SOCTS, we performed snRNA-seq of DG samples taken from young (1 month old) Tg2576 mice and wildtype (WT) littermates (n = 3/group) to identify cell type-specific alterations that could promote the hyperactivity in Tg2576 mice that occurs at this age. The number of nuclei sampled per condition (15,000) was within range of previous studies of hippocampal or DG cell types (Habib *et al*., 2016; Artegiani *et al*., 2017; Habib *et al*., 2017; Nelson *et al*., 2023). This number was sufficient to obtain proportions of nuclei for each DG cell type (see Figure 2C) that are consistent with proportions expected for mouse hippocampus (Valério-Gomes *et al*., 2018).

We obtained 6 sequencing datasets (1/mouse) containing the gene expression values for each nucleus recovered from each mouse. Prior to sorting nuclei into cell types with SOCTS, we performed initial data pre-processing steps that are required for all snRNA-seq analyses. These steps include removal of low-quality nuclei that show few detected genes (or “RNA reads”) or that show a greater than expected proportion of mitochondrial gene expression (for details of these and other pre-processing steps, see Methods). High mitochondrial gene expression is not desired because in mouse hippocampus, intact nuclei typically have a low proportion of mitochondrial gene expression (< 10; often 5% in DG; Osorio and Cai, 2021). A higher proportion can indicate that nuclei are damaged or otherwise aberrant (Osorio and Cai, 2021). Therefore, we included only nuclei that had a mitochondrial proportion of < 10% in our analyses.

As an initial validation of the pipeline, we assessed if samples were consistent with each other in the proportion of nuclei recovered for each cell type and specifically whether any significant differences were found between Tg2576 vs WT samples. The Tg2576 mouse model does not exhibit cell loss in the DG (Irizarry *et al*., 1997) and thus proportions were not expected to differ. A two-way ANOVA was performed to test if there were significant differences in Tg2576 vs wildtype (WT) mice in the proportions of nuclei found for each cell type (Figure 2C). Genotype and Cell Type were used as factors. A significant effect of Cell Type was detected (F(8, 36) = 83.73, p < 0.0001), but no effects were found for Genotype (F(1, 36) = 0.000, p > 0.9999) or the interaction (F(8, 36) = 0.2874, p = 0.9658). A Bonferroni post-hoc test with correction for multiple comparisons detected no significant differences in the mean proportions of nuclei detected in DG snRNA-seq samples from Tg2576 vs WT mice (n = 3/group; p-adj > 0.9999 for each cell type). The raw numbers of nuclei for each cell type were normalized as percentages to perform the statistics because the raw values did not show normality (Figure S3; Shapiro-Wilk: W = 0.8890, p = 0.0001; Kolmogorov-Smirnov: distance = 0.1866, p < 0.0001). The results suggest that the proportions of nuclei were similar in each genotype.

#### B. Clustering

We next performed PCA followed by UMAP clustering and present the clustering in Figure S4 where Tg2576 and WT data are positioned in adjacent plots for each panel. It is important to note that PCA and UMAP parameters are selected by the researcher and the number of clusters that emerge from a dataset is based on counting the clusters in the UMAP plot. This is a subjective process that involves iterating UMAP resolution parameters until clusters emerge with adequate space between them that they can be considered separate (Becht *et al*., 2019). A given UMAP plot can be made to display various numbers of clusters by iterating the resolution parameter to be more (approaching resolution = 1 results in more clusters) or less strict (resolution = 0 leads to few and may merge clusters). Resulting clusters in each iteration are verified to ensure each reflects an expected cell type by checking for expression of positive and/or negative marker genes known to be expressed by the cell type. Clusters are then annotated with the names that correspond to the cell types. There is no consensus regarding how many markers are necessary to identify a given cell type because the specificity of a marker can vary across biological contexts (Fischer and Gillis, 2021). Notably, it is possible to generate “clean” (no marker gene set overlaps) clusters *ad hoc* by limiting the sets of marker genes used for annotation to consist only of some but not all markers that are consistent in expression. In summary, UMAP resolution settings are often selected “by eye”, guided by user-selected marker genes, and therefore require attention to possible subjectivity (Kiselev *et al*., 2019; Chari and Pachter, 2023).

On the other hand, aspects of this process allow flexibility in clustering results which can be advantageous. Figure S4 provides an example as demonstrated in UMAP plots for each DG cell type. A PCA was performed and the top 15 PCs were used to generate each UMAP plot. We then iterated the UMAP resolution parameter for each of the cell types to be set to the minimum resolution that is needed to generate 2 UMAP clusters. For MCs, we instead selected a UMAP resolution to detect 3 clusters to show that the parameters can be altered to suit the number of clusters that are apparent when viewing a given UMAP plot. We did not perform an analysis of gene expression differences between genotypes in individual clusters found within each cell type because our goal was to identify differentially expressed genes (DEGs) in Tg2576 vs WT mice.

#### C. Benchmarking against existing methods to validate the pipeline

Our next objective was to confirm that our SOCTS-based approach can compete with a standard approach such as UMAP-based sorting. Again, the distinction is that our approach uses cell type markers to identify cell type prior to clustering instead of afterward. To make the comparison, we used 3 key performance metrics. First we determined the number of nuclei detected. Next we defined the proportion of nuclei detected using UMAP-based sorting that did not match the marker gene sets used by SOCTS. Finally, we identified the number of DEGs detected by each method for each cell type. We prioritized these metrics because they reflect, respectively, the sensitivity of nuclei detection, the specificity of nuclei identification, and the ability to identify DEGs between experimental approaches. For the UMAP-based sorting we used data from a recent study (Nelson *et al*., 2023; Figure 3A) and reanalyzed the data using SOCTS. The published study was a useful comparison because it uses a standard UMAP clustering to sort hippocampal cell types prior to marker gene labeling (referred to as a “Standard” approach below and in Figures 3 and S5-6). The Nelson *et al*. (2023) study was performed to identify transcriptional alterations in various hippocampal cell types induced after electroconvulsive shock (ECS vs Sham). Briefly, Nelson *et al*. (2023) performed ECS or Sham procedures on 2 mice per condition. Whole hippocampi were dissected from each mouse and nuclei were isolated. Nuclei were then fluorescently tagged with antibodies to the neuronal nuclear antigen NeuN. Nuclei were purified via flow cytometry to keep NeuN-positive (NeuN+) nuclei and remove non-neuronal (NeuN-) nuclei (Figure 3A).

**Figure 3.**
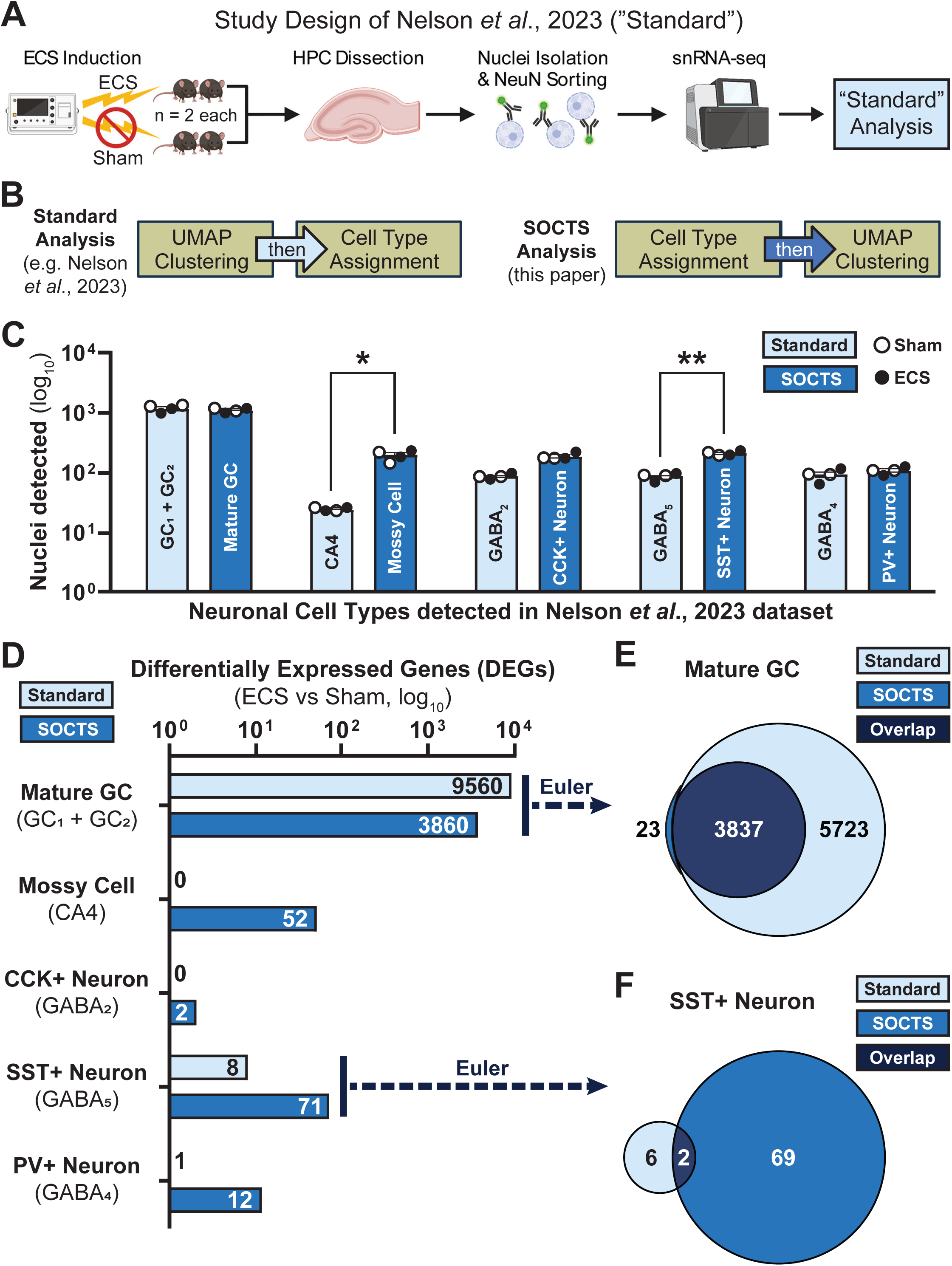
Analysis of snRNA-seq data with SOCTS may enhance sensitivity of nuclei detection and differential expression analyses vs standard approaches. We used a hippocampal snRNA-seq dataset (from the repository of the original data in Nelson *et al*. (2023); referred to as “Standard method”) to compare how our pipeline performs against standard methods when analyzing the same raw snRNA-sequencing files. **(A).** This panel shows an adapted version of Figure 1A from Nelson *et al*. (2023) that depicts their study design. Their snRNA-seq was performed on NeuN-expressing (neuronal) nuclei isolated from mice that had received electroconvulsive shock (ECS) or a sham treatment (n = 2/group). The authors reported the number of nuclei corresponding to expected cell types in the hippocampus (HPC) and performed analyses to identify DEGs (ECS vs Sham). **(B).** Diagrams are shown to illustrate the main distinction between our pipeline’s approach (SOCTS, *blue*) to cell type sorting with clustering after sorting vs the standard approach that instead begins with clustering (Standard, *light blue*). **(C).** The mean number of nuclei detected in all 4 samples (ECS & Sham) for each cell type are shown for SOCTS and the Standard (Nelson *et al*. (2023) method. The neuronal cell types defined by SOCTS (our study) are compared to the cell types associated with clusters from Nelson *et al*. (2023). A two-way ANOVA was performed to identify significant differences using the Analysis Method (SOCTS vs Standard) and the Cell Type as the factors. We found a significant effect for the Method (F(1, 30) = 11.52, p = 0.0020), the Cell Type (F(4, 30) = 451.9, p < 0.0001), and their interaction (F(4, 30) = 5.274, p = 0.0024), A Bonferroni post-hoc test with correction for multiple comparisons showed that the mean number of nuclei detected for MCs (193.0, SOCTS vs 24.5, Standard; p-adj = 0.0023) and for SST+ Neurons (208.3, SOCTS vs 86.0, Standard; p-adj = 0.0383) were significantly higher when the raw datasets were processed using our pipeline and SOCTS compared to the Standard method. No significant differences in nuclei detection were found between methods for Mature GCs (p-adj = 0.3501), CCK+ Neurons (p-adj = 0.1414), or PV+ Neurons (p-adj > 0.9999). **(D).** The number of DEGs (ECS vs Sham) detected for each cell type by each method (SOCTS vs Standard) are shown. Significance of differences for each cell type in the numbers of DEGs detected using SOCTS vs the Standard method were assessed with Fisher’s Exact Tests. The number of DEGs in Mature GCs was lower when using SOCTS compared to the Standard method (3860, SOCTS vs 9560, Standard; odds ratio (OR) = 0.2194, p < 0.0001). In contrast, the numbers of DEGs detected using SOCTS were higher compared to the Standard method in MCs (52, SOCTS vs 0, Standard; OR = infinity, p < 0.0001), SST+ Neurons (71, SOCTS vs 8, Standard; OR = 8.910, p < 0.0001), and PV+ Neurons (12, SOCTS vs 1, Standard; OR = 12.01, p = 0.0034). No significant differences were found in CCK+ Neurons in the number of DEGs found using each method (2, SOCTS vs 0, Standard; OR = infinity, p = 0.50). **(E).** The overlapping DEGs (*dark blue*) detected for Mature GCs using SOCTS (*blue*) or the Standard (*light blue*) method are shown, as well as the unique DEGs. Almost every DEG detected using SOCTS was also detected using the Standard method (3837/3760 or 99.4%), with a degree of overlap in DEGs that was significant using a Fisher’s Exact Test (OR = 192.8, p < 0.0001). **(F).** Although the number of DEGs detected in SST+ Neurons was significantly higher when using SOCTS vs the Standard method (see D), the degree of overlap between the DEGs detected via each method was significant (Fisher’s Exact Test, OR = 77.01, p = 0.0005). This may be in part due to the low number of DEGs detected using the Standard method (8 total DEGs with 2 DEG overlaps = 75%). Abbreviations: HPC, hippocampus; ECS, electroconvulsive shock; and UMAP, uniform manifold approximation and projection.

To compare the number of nuclei detected by our SOCTS-based approach vs the Standard UMAP-based approach of Nelson *et al*. (2023), we first identified which cluster labels in the data from Nelson *et al*. (2023) corresponded to the neuronal cell types in SOCTS. We pooled clusters that corresponded to the same cell type where necessary. For example, where Nelson *et al*. (2023) defined two types of GCs, we pooled those to correspond to our Mature GC cell type. We did so to allow for the most direct comparisons of each approach (SOCTS vs. Standard). Nelson *et al*. (2023) used flow cytometry to ensure that the samples of whole hippocampus contained NeuN+ nuclei, so we used SOCTS to specify the 5 neuronal cell types that we expect would be present only in the DG (GCs, MCs) or in the DG and other hippocampal subregions (CCK, SST, PV-expressing neurons). Some marker sets were adjusted. For example, *Col19a1* is present in both MCs and CA1/3 so it was not used as a marker of MCs (Cembrowski *et al*., 2016). We performed our analyses of the Nelson *et al*. (2023) samples using a modified version of our R Seurat-based pipeline (for details, see Methods). Briefly, we aligned our parameters with those of Nelson *et al*. (2023) for the pipeline operations that both approaches share. For example, the top 50 PCs were used for UMAP plots in Nelson *et al*. (2023) and thus for our reanalyses. Some processing functions in our SOCTS-based pipeline (see Methods), such as SCTransform and Harmony integration (Hafemeister and Satija, 2019; Korsunsky *et al*., 2019), were not performed in the published analysis of Nelson *et al*. (2023). Nevertheless, we decided to include these functions in the reanalysis of Nelson *et al*. (2023) because this is the current standard for snRNA-seq studies and helps reduce variability within and between samples (Hafemeister and Satija, 2019; Korsunsky *et al*., 2019; da Rocha *et al*., 2025; Hoffman *et al*., 2025).

After this reanalysis, we detected the following neuronal cell types in the Nelson *et al*. (2023) dataset using SOCTS: Mature GCs, MCs, CCK+ Neurons, SST+ Neurons, and PV+ Neurons. In the Nelson *et al*. (2023) dataset, those cell types corresponded to these cluster names, respectively: GC1 and GC2 (which we were merged), CA4, GABA2, GABA5, and GABA4. We then compared the numbers of nuclei in the data from Nelson *et al*. (2023) and our SOCTS pipeline (Figure 3C). A two-way ANOVA was performed using the Analysis Method (SOCTS vs Standard) and the Cell Type as the factors. We found a significant effect for the Method (F(1, 30) = 11.52, p = 0.0020), the Cell Type (F(4, 30) = 451.9, p < 0.0001), and their interaction (F(4, 30) = 5.274, p = 0.0024). A Bonferroni post-hoc test with correction for multiple comparisons showed that the mean number of nuclei detected for MCs was higher using SOCTS (193.0, SOCTS vs 24.5, Standard; p-adj = 0.0023) and the same was true for SST+ Neurons (208.3, SOCTS vs 86.0, Standard; p-adj = 0.0383). No significant differences were found for Mature GCs (p-adj = 0.3501), CCK+ Neurons (p-adj = 0.1414), or PV+ Neurons (p-adj > 0.9999). Therefore, SOCTS showed advantages for some cell types but not all.

We tested specificity next. Marker genes from SOCTS were used (Figure S5; Table S1) so SOCTS is represented by 100% in these comparisons. We performed this analysis on the Sham and ECS groups (n = 2 mice/group) separately to ensure treatment condition did not influence the outcome. Each panel in Figure S5 shows the proportions of nuclei labeled by SOCTS that match the cell type label (*gray*) assigned in Nelson *et al*. (2023). “Excitatory in Nelson” (*red*) refers to the excitatory neurons and “Inhibitory in Nelson” (*blue*) refers to the inhibitory neurons. We used these terms to highlight that there is disagreement between the cell type label of each cluster in Nelson *et al*. (2023) and the definition of each cell type by marker genes using SOCTS. This disagreement is reflected by the difference from 100%. Whether the disagreement was significantly different from 100% was calculated using Fisher Exact Tests. For all panels of Figure S5, the nuclei that were originally mislabeled as excitatory or inhibitory neurons were combined into one group for 2 x 2 contingency analyses. The proportions of nuclei were significantly different from 100% (i.e. SOCTS; p-adj < 0.0001). Therefore, we observed that there was an advantage in specificity using SOCTS.

We next compared the numbers and identities of DEGs (ECS vs Sham) that were detected in each cell type when using SOCTS vs the Standard analysis method (Figure 3D-F; for details, see Methods). As there were only 2 samples per condition, there are inherent statistical limitations to these comparisons. However, these limitations were mitigated by pseudobulking, the gold-standard to restrict sample size inflation (Squair *et al*., 2021; Murphy and Skene, 2022). We performed Fisher’s Exact Tests for each cell type to compare the numbers of DEGs detected using SOCTS vs the Standard analysis method (Figure 3D). The number of DEGs in Mature GCs was lower when using SOCTS compared to Standard methods (3860, SOCTS vs 9560, Standard; odds ratio (OR) = 0.2194, p < 0.0001; Figure 3D). In contrast, the numbers of DEGs detected using SOCTS were higher compared to Standard methods in MCs (52, SOCTS vs 0, Standard; OR = infinity, p < 0.0001), SST+ Neurons (71, SOCTS vs 8, Standard; OR = 8.910, p < 0.0001), and PV+ Neurons (12, SOCTS vs 1, Standard; OR = 12.01, p = 0.0034). No significant differences were found in CCK+ Neurons in the number of DEGs found using each method (2, SOCTS vs 0, Standard; OR = infinity, p = 0.50). Therefore, SOCTS shows some advantages for rare cell types, as mentioned above, but otherwise is similar to the Standard approach for more common cell types.

We next performed an analysis to determine how DEGs may differ between analysis methods, using the DEGs found in Mature GCs (Figure 3E) or SST+ Neurons (Figure 3F). For Mature GCs, almost every DEG detected using SOCTS was also detected using the Standard method (3837/3760 or 99.4%). The degree of overlap in DEGs was significant (Fisher’s Exact Test, OR = 192.8, p < 0.0001; Figure 3E). For SST+ Neurons, the degree of overlap in detected DEGs was also significant (Fisher’s Exact Test, OR = 77.01, p = 0.0005; Figure 3F). This result occurred even though the number of DEGs detected in SST+ Neurons was significantly higher when using SOCTS vs the Standard method (Figure 3D). These results provide another example of the advantages of using SOCTS.

#### D. Results of using our pipeline to address our hypotheses

##### 1. Identifying DEGs in the DG of 1 month-old Tg2576 mice that could lead to hyperexcitability

We predicted that transcriptional alterations in GCs, MCs, and possibly other DG cell types would reflect putative mechanisms by which hyperactivity could arise early in life in Tg2576 mice. To identify DEGs for specific cell types, comparisons were made between Tg2576 and WT mice at 1 month of age. Except for the use of SOCTS to identify cell types before clustering, our analysis pipeline was otherwise consistent with existing analysis practices. For example, differential gene expression was assessed in each cell type by performing pseudobulking to “pool” together samples for a given cell type into 1 pseudobulk per genotype. In practical terms, this choice sets the effective sample size for detecting DEGs by using the number of mice (n = 3) per genotype, and **not** the number of nuclei, which reduces sample size inflation and thus risk of false positives (Squair *et al*., 2021; Murphy and Skene, 2022). In line with current literature (Squair *et al*., 2021; Pan *et al*., 2023), we used Seurat’s FindMarkers tool (Hao *et al*., 2024) to perform a Poisson test (adjusted for multiple comparisons via Bonferroni’s test) to detect which genes were significantly altered between Tg2576 and WT mice in each DG cell type (Figure 4; Data S1; p-adj < 0.05).

**Figure 4.**
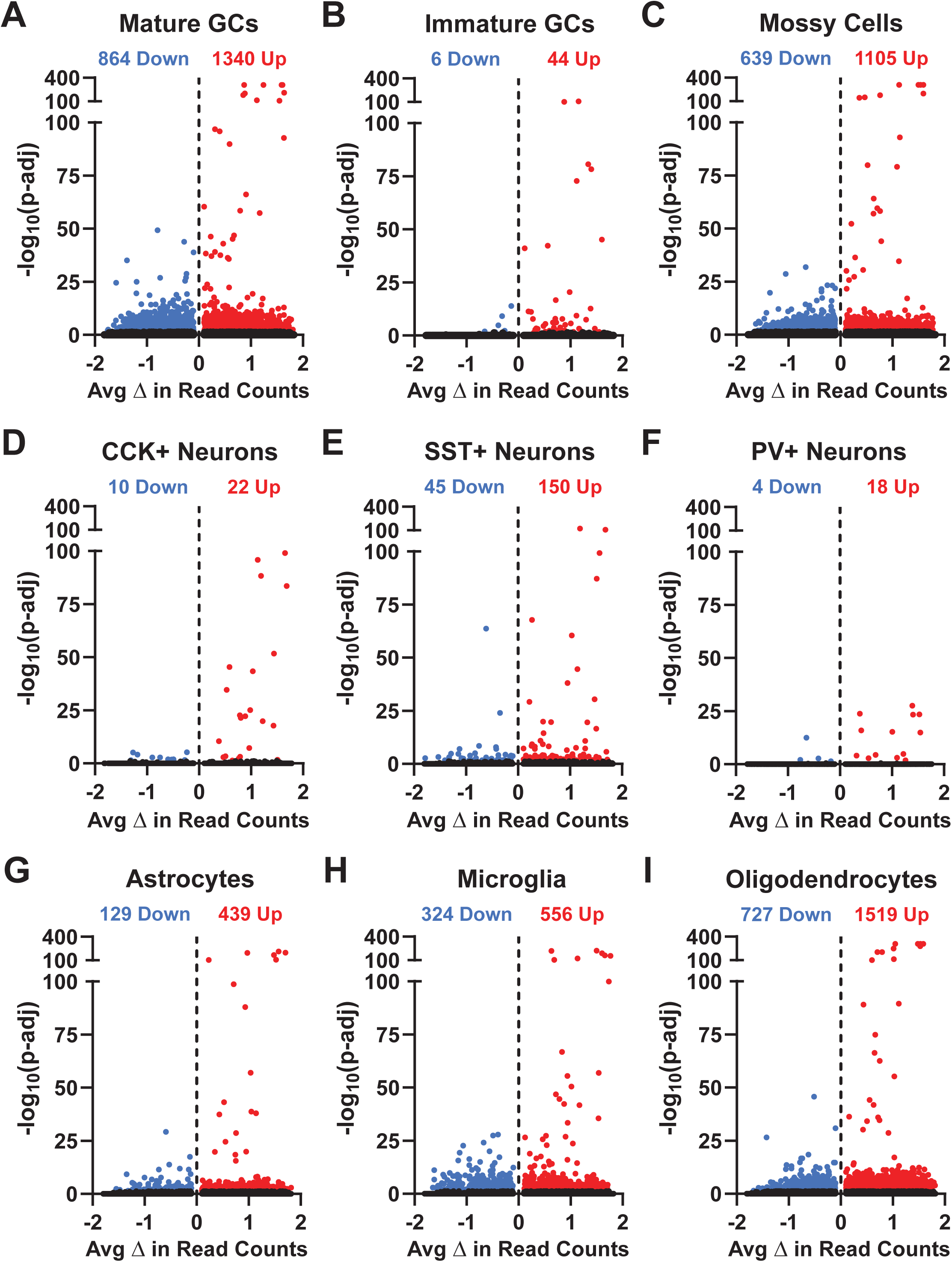
Differential gene expression between Tg2576 and WT mice can be detected for each cell type in the DG. **(A-I).** Differential gene expression analysis was performed for each DG cell type to detect DEGs between Tg2576 and WT mice (n = 3/group). Results are shown as volcano plots. The x-axis indicates the average difference in number of transcripts. The y-axis shows a negative-log transformation of the Bonferroni-adjusted p-values for the significance of difference of each gene. Individual genes = dots; *gray* dot = not significant, *red* or *blue* dot = significant increase or decrease in expression vs WT. Significant differences in expression were calculated in Seurat using a Poisson test with a Bonferroni correction (see Methods; all p-adj < 0.05). The significant or non-significant alterations in gene expression and statistics for DEGs is provided for each cell type (A-I) in Data S1.

Despite the rarity of some cell types, such as PV+ Neurons and Immature GCs, we were able to detect DEGs in each DG cell type. Indeed, we identified thousands of DEGs in total and consistent with our hypothesis that hyperexcitability depended on DEGs in GCs and MCs, we found high numbers of DEGs in these 2 cell types (Figure 4A, C). In addition, many DEGs were detected for the cell type that SOCTS suggests corresponds to Oligodendrocytes and OPCs. This was surprising but is consistent with studies showing that neuronal activity leads to altered gene expression in these cells (Gibson *et al*., 2014; Gautier *et al*., 2015). However, it should be noted that this result may also reflect the fact that this cell type had the highest number of nuclei (Figures 4I and S2). In the Discussion, we address how the altered expression of genes could regulate neuronal activity in the DG of Tg2576 mice.

Next, we performed analyses to identify common genes that were altered across DG cell types. Overall, we identified 30 genes that were changed in at least 7 of 9 DG cell types in Tg2576 vs WT mice. We found 8 DEGs increased in 9 cell types, 13 DEGs increased in 8 cell types, and 1 DEG increased in 7 cell types. There was less consistency across cell types for the DEGs that were decreased (Figure 5). Strikingly, broad alterations across the DG were in genes that regulate amyloid precursor protein (APP) processing, neurotransmission, plasticity, and mitochondrial function. The other categories of DEGs are also notable because some have already been implicated in AD or disorders of excitability (e.g. epilepsy; such as *Prnp* or *Gria1*, see Discussion). Others are novel and therefore provide new potential insights into causes of early hyperexcitability in AD.

**Figure 5.**
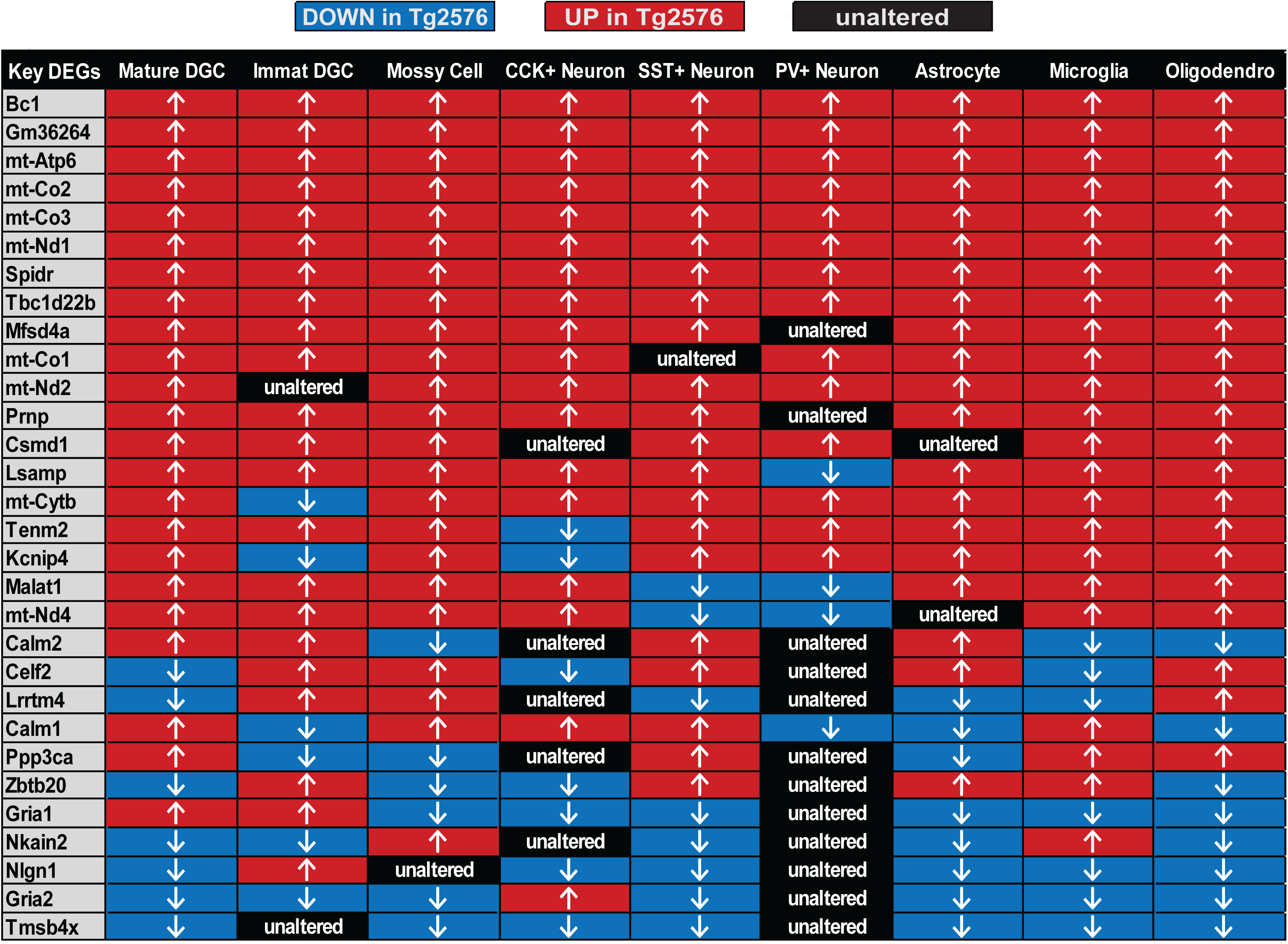
Several of the same genes show altered expression in multiple DG cell types in Tg2576 vs WT mice at 1 month of age. This panel depicts every gene that showed significant (p-adj < 0.05) increases (*red*) or decreases (*blue*) in expression in at least 7 of 9 cell types between Tg2576 and WT mice. *Black* = not significant. Many DG cell types share alterations in genes in a similar direction and related to similar functions, such as metabolism, excitability, and immune function.

##### 2. Correspondence of DEGs in mouse to human

To determine if DEGs that were altered in each cell type in Tg2576 mice are relevant to clinical AD or epilepsy, we reanalyzed a study that examined 2 public datasets of bulk proteomic alterations from AD or epilepsy patients (Leitner *et al*., 2024; Figure 6). We chose to compare our DEGs in Tg2576 mice with alterations in proteins in human AD and TLE to increase the chance of findings from mice that may translate to humans. It is acknowledged that the comparison of DEGs to proteomics and other aspects of the comparison have limitations, but we proceeded because of the opportunity to compare our data in mouse to humans, not only AD but also epilepsy.

**Figure 6.**
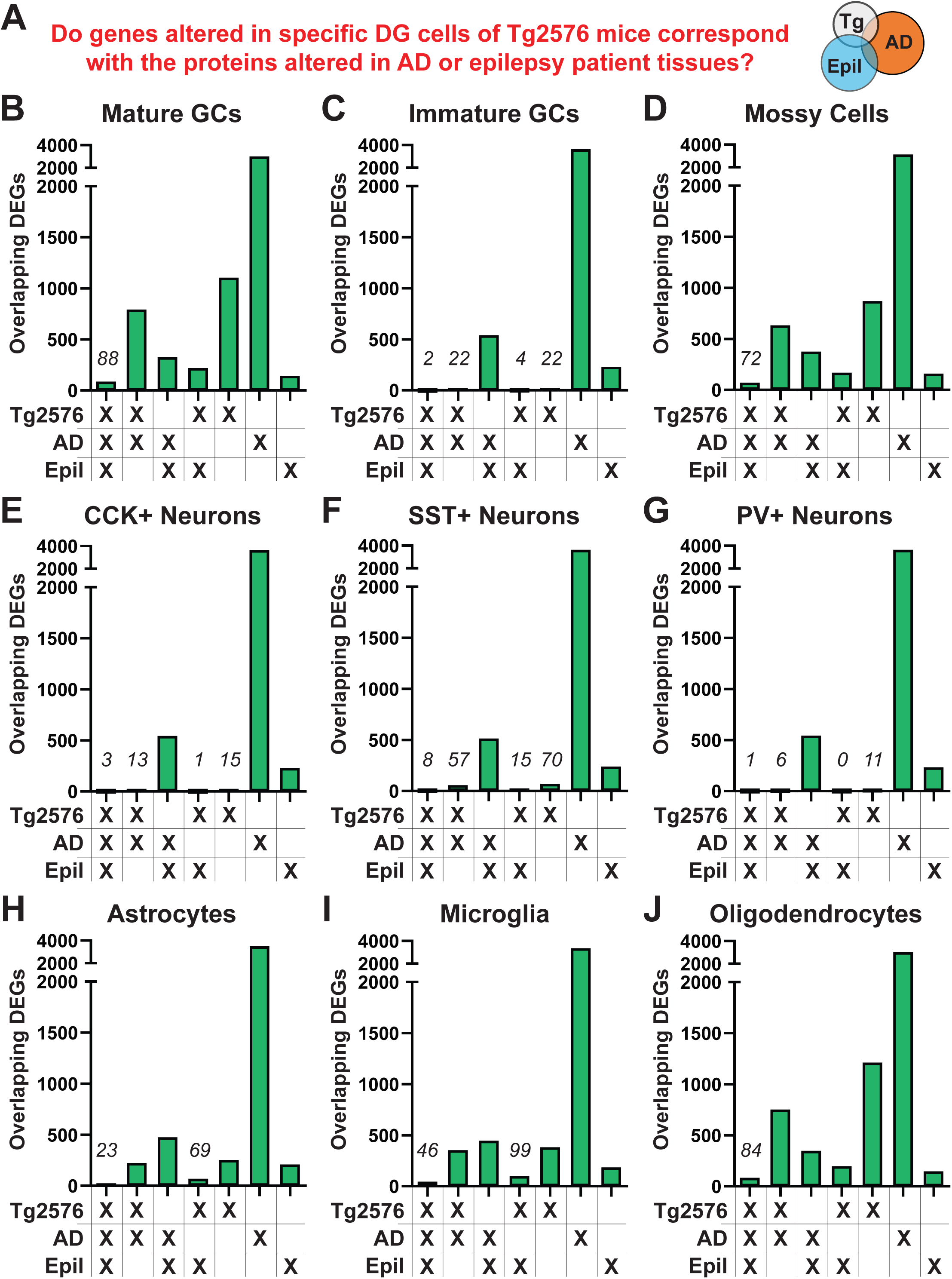
Genes altered in DG cells of Tg2576 mice share similarities with proteins altered in brains of patients with AD or epilepsy. **(A).** To understand how our results could generalize to human disease, we performed an analysis to identify the DEGs in DG cells of Tg2576 mice that overlap with the proteins altered in the brains of patients with AD or epilepsy. To perform this analysis, we compared the DEGs for each DG cell type with protein alterations derived from a published proteomics analysis (Leitner *et al.,* 2024). The Leitner *et al*. (2024) study identified overlaps in protein alterations in tissue from patients diagnosed with either AD or epilepsy compared to their respective controls. For AD, Leitner *et al*. (2024) used proteomics of several brain areas based on the NeuroPro database; which compares advanced AD to controls. For epilepsy, the authors used a hippocampal proteomics dataset from Pires *et al*. (2021) which compared protein levels altered in epilepsy vs control tissue. We performed our analysis using the data in Leitner *et al*. (2024) by overlapping our DEG lists with their lists of protein alterations and plotted the overlaps as upset plots **(B-J).** Upset plots show the intersections of DEGs and proteins altered in Tg2576 mice, human AD, and human epilepsy. The y-axes show the number of overlapping DEGs and proteins for the intersections. Data S2 provides a table for each DG cell type.

The epilepsy proteomics dataset (Pires *et al*., 2021) analyzed in Leitner *et al*. (2024) shows 777 proteins that were altered in hippocampal homogenates from 13 epilepsy patients vs 14 non-epileptic controls (see Methods, Pires *et al*., 2021, and Leitner *et al.,* 2024). The Leitner *et al*. (2024) study compared this list of 777 protein alterations in epilepsy patient hippocampus with another list of 4743 proteins altered in several brain areas of patients with AD (689/777 or 89% overlapped with epilepsy). This list of 4743 proteins altered in AD patient brain tissue corresponds to all the alterations found in patients with advanced AD, which was derived from the NeuroPro database. NeuroPro contains a meta-analysis of proteins in 13 brain regions of AD patients that show consistent alterations within the 38 proteomics datasets. For our own reanalysis, we filtered this list to remove alterations that were inconsistent in direction in different studies in the NeuroPro proteomics database. Filtering yielded 4201 consistent protein alterations for AD.

We calculated the overlap of 3 lists: (1) the 4201 proteins altered in the brains of AD patients, (2) the 777 proteins altered in the hippocampus of epilepsy patients, and (3) the DEGs in each DG cell type in Tg2576 vs WT mice (Figure 6A). Figure 6B-J depicts these overlaps as upset plots. Despite differences in age, species, diagnosis, molecule, and methodology, several genes that were altered across all DG cell types in Tg2576 mice corresponded to alterations in AD in the proteins the genes encode (Figure 6; Data S2). Strikingly, we detected a subset of DEGs in each DG cell type that overlapped with alterations in epilepsy patients, and moreover, for all 3 conditions (DEGs in Tg2576 mice, proteins in AD, proteins in epilepsy; Figure 6B-J).

##### 3. Utilizing data to identify transcription factor networks relevant to disease

Excitability can be controlled by transcription factors (TFs) that act on multiple target genes, creating a transcription factor “network.” We therefore asked if TF networks existed in our data. To do so we used a program that can highlight TF networks called “MAGIC” (Mining Algorithm For Genetic Controllers). MAGIC is used to identify which TFs have the highest proportions of target genes that are enriched in a given list of DEGs (Roopra, 2020). The target genes that are bound at genomic loci by each TF were derived from the ENCODE database (see Methods). We used MAGIC to identify putative TFs that have been shown to regulate DEGs that were altered in Mature GCs of Tg2576 vs WT mice. We performed this analysis to first assess trends in TF regulation of all 2204 DEGs altered in Mature GCs in Tg2576 mice (Figure 7A). We then filtered our analysis to identify only the top-ranked TFs that have strong influence over the DEGs that encode proteins for ion channels, transporters, or their accessory proteins and neurotransmitter receptors, since these proteins are likely to be involved in excitability (Figure 7B). In Mature GCs, our filtered lists of DEGs contained 33 up- and 31 downregulated DEGs. We found several TFs with high enrichment scores for target genes corresponding to the DEGs in Mature GCs of Tg2576 mice (Figure 7B; all results in Table S2). These TFs included several which already have been shown to play a role in hyperactivity in AD and/or epilepsy such as REST, SMARCA4, EZH2, and STAT3 (Lund *et al*., 2008; Venugopal *et al*., 2012; Grabenstatter *et al*., 2014; Hixson *et al*., 2019; Khan *et al*., 2019; Reichenbach *et al*., 2019; Choi *et al*., 2020; Mitrakos *et al*., 2020; Aron *et al*., 2023; Natali *et al*., 2023; Tipton *et al*., 2023; Hall *et al*., 2024; Al-kuraishy *et al*., 2025; Hoffman *et al*., 2025; Lagartos-Donate *et al*., 2026).

**Figure 7.**
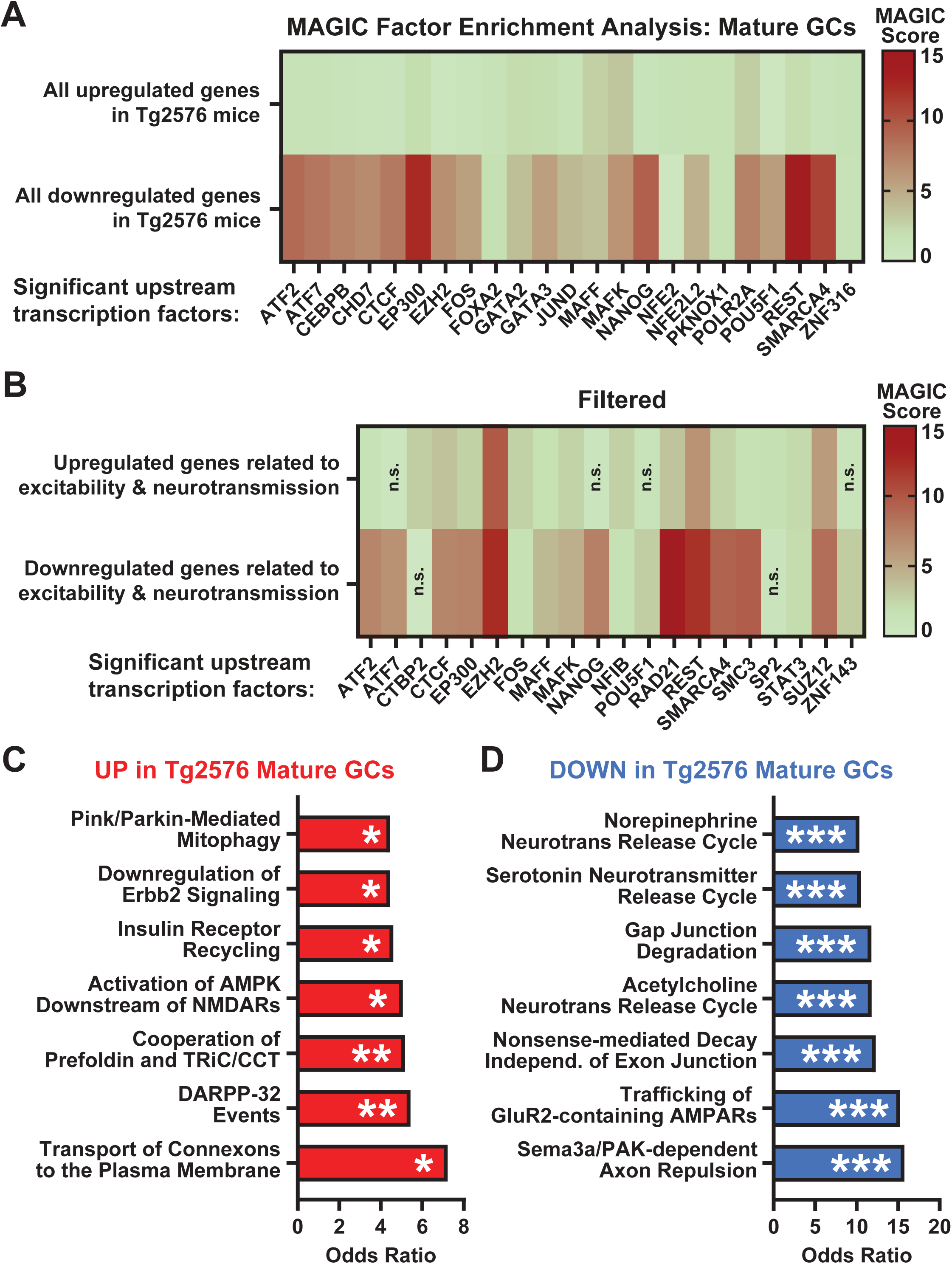
The expression of genes that can control excitability could be regulated by specific transcription factors in Mature GCs of Tg2576 mice. **(A).** MAGIC analysis was performed to identify transcription factors (TFs) that show high enrichment of binding to the DEGs identified in Tg2576 mice (see Methods and Roopra, 2020). We first performed MAGIC on the 1340 upregulated and 864 downregulated genes in Mature GCs of Tg2675 mice. Nonparametric Kolmogorov-Smirnov tests corrected with Benjamini-Hochberg FDR were performed on each DEG list to calculate which TFs showed significant enrichment (e.g. high overlap of TF target genes with DEGs). A MAGIC Score was calculated in which TFs were ranked by corrected p-values with an adjustment that selects for TFs with the most robust binding signatures. The color scale shows a range of MAGIC scores (higher, *red*; lower, *green*). All TFs shown in panel A are statistically significant (MAGIC FDR < 0.05) for enrichment of binding sites at DEGs, regardless of MAGIC scores. For both up- and downregulated DEG lists in panel A, we found several TFs with significant enrichment of binding at genomic loci of DEGs in Mature GCs of Tg2576 mice. **(B).** The DEG lists in panel A were filtered to retain only genes that encode ion channels, transporters, their accessory proteins, and receptors for neurotransmitters. We did this to identify which TFs may be most relevant to regulation of genes that could control excitability. After filtering, the DEG lists used for MAGIC analysis in panel B contained 33 upregulated and 31 downregulated genes. All TFs shown in panel B are significantly enriched (FDR < 0.05) in the MAGIC analysis. If a TF MAGIC score did not reach significance, it is marked in panel B with “n.s.” for “not significant”. **(C-D).** We performed an Ontomancer gene ontology analysis using a standard database (REACTOME; see Methods) to understand the functions that may be regulated by the 1340 upregulated (C) and 864 downregulated (D) DEGs in Mature GCs of Tg2576 mice. Like many ontology methods, Ontomancer uses Fisher Exact Tests (corrected with Benjamini-Hochberg FDR) to calculate the significance of statistical enrichment between the genes that correspond to a given biological function and the given list of DEGs. Odds ratios that correspond to the strength of enrichment for each significant biological function are shown on the x-axis. The y-axis lists the ontology categories (known as “terms”) that provide examples of the functions that are related to either up- or downregulated DEGs in Mature GCs. For significance shown on the bars: FDR = ***0.001, **0.01, or *0.05. Abbreviations: Erbb2, erb-b2 receptor tyrosine kinase 2; AMPK, 5’ AMP-activated protein kinase; NMDAR, N-methyl-D-aspartate receptor; TriC, T-complex protein Ring Complex; CCT, Chaperonin Containing TCP-1; DARPP-32, Dopamine- and cAMP-regulated phosphoprotein-32 kDa; neurotrans, neurotransmitter; GluR2, glutamate receptor 2; AMPAR, α-Amino-3-hydroxy-5-methyl-4-isoxazolepropionic acid receptor; Sema3a, semaphorin 3a; and PAK, p21-activated kinase.

##### 4. Functional analyses

To gain broader insights into the putative biological functions of genes altered in Tg2576 mice, we performed functional enrichment analyses (Subramanian *et al*., 2005). We focused on DEGs in Mature GCs given our hypotheses that Mature GCs are important to hyperexcitability early in Tg2576 mice and therefore possibly AD. However, other cell types are also relevant to AD and can be analyzed the same way (Darvas *et al*., 2026). We used the program Ontomancer (Hoffman *et al*., 2025) to perform gene set enrichment analysis (GSEA) on the list of 2204 DEGs we detected in Mature GC DEG list using the REACTOME gene ontology database (Table S2). GSEA provides an understanding of genes that are most enriched, and the REACTOME database is a common database that assigns biological functions to genes (Milacic *et al*., 2024).

For the 1340 genes *<u>increased</u>* in Mature GCs in Tg2576 mice, we found several functional pathways, some of which were surprising. One surprise was metabolism, including insulin signaling and mitophagy. This was unexpected because it would be likely to affect excitability only indirectly rather than directly. Similarly, proteostasis was notable. However, chemical and electrical synaptic transmission was prominent, which would have a direct effect on excitability (Figure 7C). For the 864 genes *<u>decreased</u>*in Mature GCs, we found multiple pathways related to chemical and electrical synaptic transmission (Figure 7D). Together the data provided the insight that defects in synaptic transmission may lead to hyperactivity that in turn causes an increase in expression of genes to control metabolism. However, other interpretations are possible as well. For example, heightened metabolism or proteostasis could potentially initiate a response of neurons to increase their activity.

## DISCUSSION

### I. Comparing our snRNA-seq pipeline to existing approaches

Our pipeline introduces several innovations to existing standard snRNA-seq methods. These innovations apply to the physical steps needed to generate the snRNA-seq dataset *<u>and</u>* to its analysis. First, we minimize the time needed to remove the DG and maximize tissue integrity. We also incorporate PIP-seq to keep nuclei stable to decrease the need for long processing time and allow more time for transport. While PIP-seq itself is not novel, implementation of PIP-seq involved a new modification so it could be used with a common, user-friendly kit (10X Genomics). By combining these innovations, our pipeline improves upon existing snRNA-seq methods.

The analytic innovation of our approach is to separate cell types based on their marker genes *<u>prior</u>* to clustering. In contrast, existing snRNA-seq approaches separate cell types *<u>after</u>* using clustering. Both approaches define cell type by marker gene expression. However, our approach uses this definition of cell type to both sort our nuclei and interpret our results, whereas the common approach uses this definition of cell type primarily to interpret results but not to sort nuclei.

It may be useful to describe the existing snRNA-seq analysis approach in more detail to understand the advantages and disadvantages of our approach. The existing approach performs the initial sorting of cell types using a mathematical definition of cell type based on the separation of the clusters after PCA and subsequent UMAP visualization. Afterwards, cluster identities are verified using marker genes to define different cell types. If those clusters indeed contain nuclei with expected markers, they are said to be validated. However, clusters may have a combination of nuclei of different types, making it hard to separate cell types (Figures 3 and S5, and see Yao *et al*., 2024). That is why we chose to specify cell types as the initial step. A potential limitation with our approach is that specifying cell types before clustering does not allow for discovery of new cell types. However, this may be an advantage to answer questions that are related to the common view of cell types.

In summary, our approach is valuable when ambiguous clustering occurs or cannot be ruled out. In addition, potential sources of error in cell type definition are prevented from influencing DEG detection. Existing methods are less problematic when clusters are clear and there are multiple well-studied positive and negative marker genes for cell types.

Other research groups have also introduced software programs for snRNA-seq analysis that allow users to identify cell type without using clustering to perform the sorting step. Those approaches, however, differ from our SOCTS analysis in that they are either not applicable to mouse DG, derive *de novo* marker gene signatures from the user dataset to compute putative cell types, or use large marker gene reference datasets. These reference datasets in practice can be difficult to reproduce, perhaps in part because they depend on degrees of expression (not presence/absence) and can often involve many marker genes that have not been validated in the tissue to be studied (Aran *et al*., 2019; Zhang *et al*., 2019; Fischer and Gillis, 2021; Hao *et al*., 2021; Domínguez Conde *et al*., 2022; Ianevski *et al*., 2022). SOCTS analysis avoids these issues by maintaining the same biological definition of cell type for sorting and interpretation. It does so by using the presence and/or absence of markers that have been validated by other groups in the mouse DG literature.

Our benchmarking experiments (Figures 3 and S5) help demonstrate how these differences in cell type identification can impact snRNA-seq results and highlight additional advantages of our approach. The original results in Nelson *et al*. (2023) were obtained using standard approaches where cell types were defined after clustering. In our SOCTS-based reanalysis of this dataset, where cell types were defined before clustering, we found some similarities and several differences from Nelson *et al*. (2023). We detected similar proportions of nuclei for GCs (Figure 3C) and we found most DEGs in GCs were similar when SOCTS was compared to their method (Figure 3E). This is likely to be due to the high degree of cell type agreement (93-94%; Figure S4) in the GC nuclei that were found via SOCTS or Nelson *et al*. (2023). However, it should be noted that fewer DEGs were found overall (3860 vs 9560; Figure 3D) using our approach for DEG detection vs the approach used in Nelson *et al*. (2023). This may be because our approach has improved preprocessing steps and more conservative parameters for detection of differential expression, which was done to minimize Type I errors (Hafemeister and Satija, 2019; Squair *et al*., 2021). However, our overall analysis pipeline may nevertheless provide enhanced sensitivity in detection of bona fide DEGs. This could be due to greater statistical power that arises due to the ability of SOCTS to detect more nuclei per cell type, as shown in the MCs and SST+ Neurons (Figure 3C). Statistical power would be further enhanced by the ability of SOCTS to reduce the heterogeneity (and thus population variability) between nuclei that are assigned to a cell type. This effect is demonstrated via the enhanced cell type specificity for SOCTS vs Nelson *et al*. (2023) in Figure S5 and could be responsible for the enhanced DEG counts found for all cell types except CCK+ Neurons in Figure 3D.

It should be noted that comparisons to other studies could be made. However, Nelson *et al*. (2023) is both representative of many current methods used in snRNA-seq analysis and readily demonstrates that SOCTS can be used even for smaller scale studies (n = 2-4). However, although SOCTS can be used with a small number of samples, a larger number of mice is recommended to ensure sufficient sampling and statistical power (Halasz *et al*., 2025).

### II. Implications of the outcomes for our hypotheses

#### A. DG cells showed many alterations in gene expression by 1 month of age in Tg2576 vs WT mice

We had predicted that Mature GCs and MCs in Tg2576 mice would show many alterations in gene expression at 1 month of age. We made this prediction because those cells show changes in excitability at this age (Alcantara-Gonzalez *et al*., 2021; Alcantara-Gonzalez *et al*., 2025). In addition, there are IIS in vivo at this age and their origin appears to be the DG (Lisgaras and Scharfman, 2023b). In support of this idea, we detected many DEGs in Mature GCs (864 down/1340 up) and MCs (639 down/1105 up). However, neuronal activity in the DG is influenced by the other DG cell types as well (Scharfman, 2011; Scharfman, 2019). It was therefore expected that we would detect DEGs in other DG cell types and we did (Figure 4). Overall, these widespread alterations in gene expression support the hypothesis that early dysregulation of gene expression in the DG could be a mechanism by which the DG can initiate IIS. Although this is a mouse model and not clinical AD, finding similarities in the model and clinical AD (see below) suggests that the model predicts aspects of AD pathophysiology in humans.

#### B. Several genes were altered in similar directions across multiple DG cell types in Tg2576 mice

We did not expect to see the same DEGs in multiple DG cell types (Figure 5). Nevertheless, many DEGs were shared by multiple DG cell types and they were often altered in the same directions. Notably, several of these genes have been implicated in mechanisms of AD and/or other disorders with brain hyperactivity. For example, we saw that genes known to regulate the production and clearance of Aβ were increased across most or all DG cell types. The most prominent of these genes were *Bc1* (brain cytoplasmic 1) and *Prnp* (prion protein), although these genes have many additional functions related to plasticity, immune regulation, and others (Griffiths *et al*., 2012; Briz *et al*., 2017; Lathe and Darlix, 2017; Zhang *et al*., 2018; da Silva Correia *et al*., 2024). Shared DEGs were found across DG cells (*Lsamp*, *Tenm2*, *Calm1/2*, *Kcnip4*, *Mfsd4a*, *Celf2*, *Zbtb20, Gria1/2*, *Tmsb4x*, *Nlgn1*, *Ppp3ca*, and *Nkain2*) that are also involved in excitability, neurotransmission, and plasticity (Rubin *et al*., 1999; Pang *et al*., 2010; Massone *et al*., 2011; Sanz *et al*., 2015; Fang *et al*., 2016; Cid *et al*., 2021; Favaro *et al*., 2024; Gupta *et al*., 2025; Merrion *et al*., 2025; Zeng *et al*., 2025; Hua *et al*., 2026), suggesting that multiple cell types could potentially contribute to increased excitability via common pathways. We also saw shared DEGs that can regulate complement immune signaling (such as *Csmd1*) suggesting that early activation of the immune system could accompany or even promote increased excitability in Tg2576 mice (Werren *et al*., 2024). Taken together, the results provided several new insights for causes of hyperexcitability as well as surprising findings regarding other cellular pathways.

Intriguingly, we saw a shared set of mitochondrial-encoded genes that were primarily increased (*mt-Atp6*, *mt-Co1/2/3*, *mt-Nd1/2/4*, and *mt-Cytb*). One might expect snRNA-seq would exclude mitochondrial genes because mitochondria exist outside the nucleus. However, it is now known that mitochondrial RNAs can regulate nuclear gene expression and that mitochondria bind to the nuclear pore complex. Therefore, it is expected that some mitochondrial RNA would be detectable in the cell nucleus, even if it is synthesized in the extranuclear mitochondria (Muneretto *et al*., 2024; Sriram *et al*., 2024; Khatri and Blumental-Perry, 2025; Menendez-Montes *et al*., 2026). The results are important because mitochondrial activity and mt-gene expression are increased early in AD (Naia *et al*., 2023; Hung *et al*., 2025).

#### C. Mature GCs and MCs in Tg2576 mice exhibited alterations in genes that can impact hyperactivity

As mentioned above, we found alterations in both Mature GC and MCs in genes that regulate neuronal activity, excitability, neurotransmission, and plasticity. Overall, most of these DEGs fell into similar categories in Mature GCs and MCs. Before we discuss the many similarities, there is one notable difference to highlight. In MCs and not GCs, there were alterations in genes that encode voltage-gated calcium channels (*<u>down</u>*: *Cacna1c* & *Cacna2d1*; *<u>up</u>*: *Cacna1b & Cacna1d*). This difference suggests alterations in calcium channels in MCs could explain why MC intrinsic properties showed elevated excitability in 1 month old Tg2576 mice but intrinsic properties in GCs only became abnormal at 2-3 months (Alcantara-Gonzalez *et al*., 2021; Alcantara-Gonzalez *et al*., 2025). However, differences in GCs vs MCs at 1 month may also be explained by other genes altered in different directions in GCs and MCs. These genes include the ion channels *Hcn1* & *Kcnma1* (Data S1).

Regarding DEGs that were altered in the same direction in both GCs and MCs, several are implicated in excitability, such as genes that encode potassium channels and their regulator proteins (*<u>down</u>*: *Kcnab1*, *Kcnd2*, *Kcnq3,* & *Kctd16*; *<u>up</u>*: *Kcnb1*, *Kcnh1, Kcnip3/4, Kcnt2,* & *Kctd4/17*). Next, there were several shared gene alterations in glutamate receptors and their regulator proteins (*<u>down</u>*: *Gria2/3*, *Grid2*, *Grik2*, *Grin2b*, *Grip1*, & *Grm5/7; <u>up</u>*: *Grin1* & *Grina*), and in GABAergic receptors and their regulator proteins (*<u>down</u>*: *Gabrb3*; *<u>up</u>*: *Gabrap* & *Gabrb2*). Although in fewer numbers, we also saw shared alterations in genes that encode sodium channels (*<u>down</u>*: *Scn1a* & *Scn2a*; *<u>up</u>*: *Nalcn* & *Scn2b*) and ryanodine receptors (*<u>down</u>*: *Ryr2/3*). Finally, we saw a shared increase in the gene that encodes a serotonin receptor, *Htr4*, and is of interest because it is implicated in epilepsy (Leitner *et al*., 2022; Sourbron and Lagae, 2022). It should be noted that some gene alterations appear to reflect pathological mechanisms while other alterations could be compensatory, depending on the direction in which genes were altered. For example, in GCs and MCs, increases in *Nalcn* and *Scn2b* expression would be expected to increase neuronal activity. However, these alterations were accompanied by decreases in *Scn1a* and *Scn2a*, two gene alterations that would be expected to decrease neuronal activity. Thus, future functional manipulations of specific pathways will be required to dissect the impacts of individual DEG pathways on early hyperactivity in Tg2576 mice. Because we observed so many similarities in altered gene expression in GCs and MCs, the common genes may provide promising targets for future studies. For example, a future direction would be to test if upstream regulatory TFs can be identified. Below, we provide an example. Such TFs may provide druggable targets for a clinical strategy (Wang *et al*., 2025a).

#### D. Transcription factors in Tg2576 mice may regulate genes in the DG that can control excitability

In epilepsy, a critical mechanism of hyperactivity involves activation of the TF Stat3 by Janus-activated kinase 1 (*Jak1*) in Mature GCs. Inhibition of this TF pathway via a blockade or genetic reduction of Jak1 or Stat3 can induce a profound suppression of hyperactivity and a restoration of spatial memory in models of epilepsy (Lund *et al*., 2008; Grabenstatter *et al*., 2014; Hixson *et al*., 2019; Tipton *et al*., 2023; Carrel *et al*., 2025; Hoffman *et al*., 2025). Indeed, a recent study in a mouse model of epilepsy demonstrated long-lasting reductions in hyperactivity following a 2-week treatment with tofacitinib, a Jak1 inhibitor that is FDA-approved to treat some inflammatory conditions (Hoffman *et al*., 2025).

Strikingly, we detected a robust increase in *Jak1* expression in Mature GCs of 1 m.o. Tg2576 mice. In hindsight, it would be expected that *Jak1* would be altered in a model of AD because Jak1 receptor signaling is already known to exacerbate memory deficits and neuropathology in AD, but it is currently presumed that Jak1 acts in AD primarily by influencing inflammation (Chiba *et al*., 2009; Eufemi *et al*., 2015; Reichenbach *et al*., 2019; Choi *et al*., 2020; Lyra *et al*., 2021; Mehla *et al*., 2021; Al-kuraishy *et al*., 2025). Our results suggest that Jak1 signaling could also contribute to hyperactivity in early stages of AD, possibly by reducing inflammation and hyperactivity. Therefore, a promising future direction would be to test if tofacitinib could be repurposed in AD to reduce hyperactivity and cognitive deficits.

We also identified other prominent TFs that could regulate DEGs in Mature GCs in our 1 m.o. Tg2576 mice using the MAGIC analyses shown in Figure 7A-B. After focusing on encoded proteins related to excitability (ion channels, transporters, neurotransmitters and receptors), we detected TFs involved in the activity of the JAK1-STAT3 pathway: STAT3, EZH2, SUZ12, and SMARCA4 (Table S2). Remarkably, many of the putative targets of these TFs were also DEGs in Mature GCs and MCs in Tg2576 mice (such as *Grm5/7* and *Kcnq3*; Figure 5). Furthermore, many DEGs were altered in the same directions, supporting other results suggesting many DEGs showed similar directions of alteration in different DG cell types.

#### E. Genes altered in the DG of Tg2576 mice overlap with proteins altered in human AD and epilepsy

We made comparisons to clinical proteomics datasets to assess the ability of our pipeline to detect DEGs in a model of AD that can be recapitulated in patients with AD or epilepsy (Figure 6). As described in the Results, the obstacles included differences in age, species, diagnosis, neuropathology, gene vs protein expression level correlations, brain regions sampled, and use of snRNA-seq vs bulk-tissue proteomics. However, we detected overlaps for all respective cell types between the DEGs found in our snRNA-seq of Tg2576 mice and protein alterations in human AD brains (Figure 6). We also detected overlaps of protein alterations in human epilepsy with the DEGs in each DG cell type. There was an exception regarding PV+ Neurons (Figure 6G). This is likely due to lower numbers of DEGs detected in PV+ Neurons vs other cell types. In addition, there was a relatively small list of protein alterations in the human epilepsy vs AD groups compared to their controls (777 in epilepsy vs 4201 in AD). Nevertheless, the findings generally supported the use of our snRNA-seq pipeline to identify disease-relevant gene expression alterations in small brain areas like the DG.

We also addressed if the shared DEGs in Tg2576 and human AD or epilepsy patients were consistent with our hypothesis. As predicted, we found that major genes and proteins that are altered in common across all 3 conditions can regulate neurotransmission and excitability. Some examples are genes that encode metabolic and synaptic proteins, as well as K^+^ or Ca^2+^ channels. Some of the DEGs have already been studied in AD or epilepsy but others appear novel, such as *Lsamp* (limbic system-associated membrane protein) and *Camkv* (Calmodulin kinase-like vesicle-associated pseudokinase).

### III. Conclusions

Together, our results indicate that our snRNA-seq pipeline provides an advantageous, user-friendly method to investigate gene alterations in small circuits of the brain. Our sample preparation procedures introduce technical innovations that can be helpful in generating rigorous snRNA-seq datasets. Our analysis procedures demonstrate effective cell type annotation of nuclei in our snRNA-seq dataset (and in other datasets) without the need for clustering beforehand. Our annotation approach is well-suited to study additional brain regions because the marker gene panels used by SOCTS can be modified to detect additional cell types.

We demonstrated that, when we used our pipeline to test our hypotheses in an AD mouse model, we can use our approach to find numerous DEGs throughout the cell types of the DG. We identified numerous DEGs that were shared by multiple DG cell types and those that were cell type-specific. In line with our hypothesis, many of the DEGs that were altered in DG cell types were genes that could regulate brain hyperactivity. Moreover, this was true for DEGs in Mature GCs and MCs, consistent with our electrophysiological findings in our past studies of this mouse model at this age. We showed that many of the DEGs in DG cell types in our AD mouse model can be detected in proteomics datasets of brain tissue from human AD and epilepsy patients. This result helps to validate both that our DEGs detected in mouse DG are robust enough to generalize to clinical datasets, and that our DEGs in this AD mouse model related to hyperactivity may help inform therapeutic strategies to reduce hyperactivity in the clinic. Because we found druggable TF pathways that could help control excitability in the DG, our findings suggest possible avenues for drug repurposing to target these pathways in AD and other disorders with hyperactivity.

## METHODS

### I. Experimental animals

#### A. The Tg2576 mouse model of AD

In this study, we used the Tg2576 mouse model of AD and performed analyses in comparison to wildtype (WT) littermates (male only; n = 3 mice/genotype). The Tg2576 mice express transgenic human APP695 (hAPP) under control of the hamster prion promoter (Hsiao *et al*., 1996). This hAPP construct contains the Swedish mutations associated with familial AD (K670M, N671L) that increase production of Aβ from hAPP. Tg2576 mice constitutively express high levels of hAPP (Hsiao *et al*., 1996). By 1 month of age, these mice exhibit generalized IIS which persist throughout life (Kam *et al*., 2016; Lisgaras and Scharfman, 2023a, b; Chartampila *et al*., 2024). Soluble Aβ oligomers in MCs are detected at 1 month of age and deficits in novel object location and recognition tasks are detectable by 3 months but not 1 month (Duffy *et al*., 2015; Alcantara-Gonzalez *et al*., 2025). Plaque is first detected at 6 months (Hsiao *et al*., 1996) and robust after 9 months (Criscuolo *et al*., 2024). The exact ages of Tg2576 and WT mice used in this study were matched (mice in each group were respectively 38, 38, and 44 days old) and an average of 40 days old. It is acknowledged that the Tg2576 mouse model is only one of many models and all have inherent limitations (Chin, 2011; Granzotto *et al*., 2024). We chose to use 3 mice/group because a common practice is to start with this sample size, although we acknowledge that higher sample sizes could be used to ensure maximal statistical power (Cembrowski *et al*., 2016; Nelson *et al*., 2023; Halasz *et al*., 2025).

#### B. Animal care, breeding, and genotyping

All experimental procedures were approved by the Institutional Animal Care and Use Committee (IACUC) at The Nathan Kline Institute and experiments were carried out in accordance with the National Institutes of Health (NIH) guidelines. Experimental mice were housed 2-4/cage in standard cages (26 cm wide × 40 cm long × 20 cm high, Allentown) with corncob bedding (1/4” Bed-o’Cobs, #10003861, W.F. Fisher), a 2.5 g nestlet (1” x 1”, NES7200, Gusner Enterprises) in each cage, and a light:dark cycle of 12 hours. Cage changes were performed at least weekly. Mice of different sexes were housed in separate cages after weaning.

Tg2576 mice were bred in-house using the F1 hybrid of male heterozygous Tg2576 and female non-transgenic mice (C57BL6/SJL, #100012, Jackson Labs). Breeder pairs were housed with enrichment (plastic dome) and fed chow that is specialized for breeding (Purina LabDiet 5008, W.F. Fisher). Litters were weaned at 23-25 days old and fed standard rodent chow (Purina LabDiet 5001, W.F. Fisher) *ad libitum*. Genotyping of tail samples was performed using a protocol to detect expression of the APP695 gene (Transnetyx). Analyses were blind to genotype until after detection of DEGs in each cell type.

### II. Data acquisition

#### A. Day 1. Tissue harvesting

##### 1. Brain harvesting and slice preparation

As shown in Figure S1, mice were anesthetized by inhalation of 1-2% isoflurane (Piramal, #0010250P, Patterson Veterinary Supply). Mice then received a terminal dose of intraperitoneal injection of urethane (2.5 g/kg, #94300, Sigma) and were placed on ice. After reaching a surgical level of anesthesia, mice were perfused transcardially with a chilled (4°C), sucrose-based version of artificial cerebrospinal fluid (aCSF) containing a low concentration of Ca^2+^ and high Mg^2+^ to reduce synaptic transmission (in mM; Sigma-Aldrich product numbers are shown): 90.0 sucrose (#S8501), 2.50 KCl (#P9541), 1.25 NaH_2_PO_4_ (#S5011), 4.50 MgSO_4_ (#M1880), 25.0 NaHCO_3_ (#S5761), 10.0 D-glucose (#G8270), 80.0 NaCl (#S9888), and 0.5 CaCl_2_ (#C5080; pH 7.4). It was aerated with carbogen (95% O_2_ and 5% CO_2_; All-Weld Products) for 15 min prior to addition of CaCl_2_ and then after another 15 min of aeration before use. In our past studies, these conditions were ideal for preparing slices for electrophysiological recordings (Alcantara-Gonzalez *et al*., 2021; Alcantara-Gonzalez *et al*., 2025).

After perfusion, brains were removed and immersed briefly in cold sucrose-aCSF in a beaker surrounded by ice. Brains were patted dry with a Kimwipe. The cerebellum and olfactory bulbs were removed with a razor blade. The dorsal surface of the brain was mounted onto a Vibratome stage with cyanoacrylate glue to allow slicing of horizontal sections starting at the ventral surface. Once mounted, the stage was filled with chilled sucrose-aCSF and kept cold during slicing by periodically placing frozen sucrose-aCSF cubes into the stage. A Vibratome (#VT1200S, Leica) was then used to quickly prepare bilateral horizontal slices at a thickness of 500 µm. Slices were placed in a sterile petri dish (Corning Cell Culture Dish 35 mm × 10 mm, #430165, Fisher) filled with cold sucrose-aCSF in an ice bucket. Once all slices containing DG were collected, the petri dish and slices were placed on ice in a shallow insulated box that fits on the stage of a low-magnification dissection microscope (Nikon SMZ800, Micro-Optics Precision Instruments). It should be noted that a potential limitation to this process is difficulty sampling from the most septal part of the hippocampus. However, other slice orientations that allow such sampling, such as coronal or transverse, can be used if desired.

##### 2. Dissection to obtain DG tissue samples

A container containing dry ice and labeled sample tubes was placed next to the dissection area. The tubes were the 1.5 mL “Sample Dissociation Tubes” that are included in the 10x Genomics Nuclei Isolation Kit (see below). Placing DG tissue into these tubes for freezing and storage enabled the samples to remain in the same tube when thawed at the beginning of nuclei isolation and homogenized (see below).

For each hemisphere of a slice, a biopsy punch (1 mm-diameter, #15110-10, Ted Pella) was used to rapidly remove the DG. While the punch contained some of the tail end of the CA3c cell layer, procedures later could filter out CA3 neurons by their markers. Tissues were immediately flash-frozen on dry ice as described above. The lid of the tube was kept open during tissue transfer, and therefore, the lid of the dry ice container was closed when possible to minimize condensation inside the tube. Once all DG tissue was removed (typically from 6 slices per mouse), the Sample Dissociation Tubes containing DG were stored at −80°C until processing.

#### B. Day 2. Isolating and assessing DG cell nuclei

##### 1. Dissociation of tissue and isolation of nuclei

We followed a modified protocol derived from the Sample Prep User Guide for the 10x Genomics Chromium Nuclei Isolation Kit (#PN-1000494). The Debris Removal Buffer was not used and was replaced by a wash step as advised by Fluent technicians. Also as advised, the Nuclei Wash and Suspension Buffer in this kit was replaced with a buffer that is compatible with PIP-seq made by Fluent (6X Nuclei Suspension Buffer, #FB0004726). Fluent Nuclei Suspension Buffer was prepared as advised as described below.

We began Day 2 by preparing the following solutions on ice (volumes provide enough solution for at least 6 mice as described above). To make the Lysis Buffer, we combined 6600 µl of the Lysis Reagent, 6.6 µl Reducing Agent B, 66 µl Surfactant A, 168.5 µl 40X RNAse Inhibitor (all from the 10x Genomics Chromium kit), and 66.73 µl 100X Halt protease and phosphatase inhibitor (#78440, Fisher). To make the Fluent Nuclei Suspension Buffer, we combined 2171 µl 6X Nuclei Suspension Buffer (Fluent), 520 µl 25% bovine serum albumin (#FB0004728, Fluent), 325 µl 40X RNAse Inhibitor (from 10x Genomics Chromium kit), 130 µl 100X Halt inhibitor (Fisher), and 9854 µl purified water. All work surfaces, gloves, and instruments were decontaminated to remove RNAses using RNAse Zap (#R2020, Sigma). RNAse-free disposables were used.

To process each DG sample, the Sample Dissociation Tube was removed from dry ice and 200 µl of Lysis Buffer was added. The tissue was then homogenized until a homogeneous slurry formed using a pestle from the 10x Genomics Chromium kit mentioned above. Once all samples were dissociated, 300 µl Lysis Buffer was mixed into each sample via pipette trituration (5 cycles) and samples were placed on ice to incubate for 8 minutes.

##### 2. Purification of isolated nuclei

To liberate nuclei from lysed cells, each sample was triturated 15 times and immediately dispensed into the Nuclei Isolation Column that is inside each Sample Collection Tube (both from the 10x Genomics Chromium kit, placed on ice). We centrifuged the samples in a 4°C microcentrifuge (Sorvall Legend Micro R21, #75002490, Fisher) for 20 seconds at 16,000 relative centrifugal force (rcf). This step is performed to trap debris in the column and away from the nuclei suspension that flows into the Sample Collection Tube. The columns were discarded and each sample tube was vortexed at maximum speed for 10 seconds to resuspend nuclei. Each sample was then centrifuged for 3 minutes at 500 rcf to form a nuclei pellet. The supernatants were gently removed with a 200 µl pipette until roughly 200 µl remained (to help preserve pellet). For the first wash step, each pellet was resuspended by adding 800 µl Nuclei Suspension Buffer and triturating 15 times with a 1 ml pipette tip. Samples were then centrifuged for 5 minutes at 500 rcf. Supernatants were removed as before, leaving ∼200 µl behind with the pellet. Pellets were then resuspended for the second wash. The second wash was performed exactly like the first wash, except that the supernatant was removed from the second wash until only enough remained to cover the pellet. After the supernatant was mostly removed, each pellet was resuspended in a volume of no more than 50 µl of the Nuclei Suspension Buffer using a 200 µl pipette tip to mix 5 times. All samples were maintained on ice for the next steps, which should be performed immediately to preserve quality.

##### 3. Assessment of quality and concentration of nuclei

We used a brightfield microscope (Model BX-51, Olympus of America) digital camera (Lumenara 3.0, Teledyne) and software (InfinityAnalyze, Teledyne) at 16X magnification to assess the quality of the nuclei we obtained and to quantify the concentrations of nuclei in each sample. To do this, each sample was resuspended (rapid trituration with 20 µl tip 15 times) and a 20 µl solution was prepared that contained 2 µl of the nuclei suspension, 9 µl of 0.4% trypan blue (#15250-061, Fisher), and 9 µl Nuclei Suspension Buffer (Fluent). This 1:10 solution was triturated 15 times and 10 µl was dispensed into a disposable hemocytometer (#50-131-1352, Fisher) under illumination. Nuclei of each sample were considered acceptable if samples showed minimal or no clumping of nuclei into aggregates, cellular debris, and dead (punctate) or shrunken (pyknotic) nuclear bodies.

Nuclei concentrations were estimated using hemocytometer instructions to obtain nuclei/µl values (average nuclei counted in 5 grid chambers, multiplied by 10 to account for the dilution). These nuclei concentration estimates were used to ensure that no more than 5000 nuclei were contained in the 4 µl suspension (1250 nuclei/µl) that would be loaded into the PIP tube for later processing. The ideal capacity of the PIP-seq version we selected (T2 PIP-seq v4.0PLUS, see below) is 5000 nuclei per sample and overloading is not recommended by Fluent. Later steps described below were used to standardize numbers and transcriptional quality of nuclei.

#### C. Day 2. Preparing nuclei samples for PIP-sequencing

Once visual quality and concentration of nuclei were assessed, we followed the exact protocol in the User Guide for the PIP-seq T2 v4.0PLUS 3’ Single Cell RNA kit (#FBS-SCR-T2-8-V4.05, Fluent). In that protocol, all steps from 1-18 (p. 20-24; document FB0001026, version 12.9) can be performed in the user’s own lab. These steps require a digital “dry bath” (#FBS-SCR-PDB; Fluent) and a vortex with an attachment that can rotate 90 degrees to allow horizontal and vertical mixing (#FBS-SCR-DVM; Fluent). Both instruments can fit in a backpack if transport is necessary. The Fluent instruments were loaned to us by the sequencing core to allow us to prepare and stabilize PIP-seq samples for submission to the sequencing core. Loaning instruments might also be possible at other sequencing cores and would have similar advantages. Another digital heat block or a different vortexer can also be used instead of those from Fluent. In the current study, instruments were from Fluent.

##### 1. Encapsulation of nuclei in PIP emulsions

To briefly describe the steps from the PIP-seq User Guide (e.g. Section 5.1 Capture and Lysis), we prepared the 0.5 ml PIP tubes (#FB0003913, Fluent) by allowing them to thaw on ice while the dry bath was preheated to 66°C. Nuclei were resuspended and 4 µl of nuclei suspension was added to each respective PIP tube. To maximize RNA integrity, 0.5 µl of Protector RNAse Inhibitor (40 Units/µl, #3335399001, Sigma) was added to each PIP tube prior to adding the nuclei suspension. Each PIP tube was mixed via gentle trituration (10 cycles with 200 µl tip) and kept at room temperature for subsequent steps. The Partitioning Reagent (280 µl; #FB0001550, Fluent) was added to each tube. The PIP tubes were vortexed as described in the protocol in the User Guide to encapsulate single nuclei into individual partitions of PIP emulsion. The tubes were allowed to stabilize so that the emulsion layer separated from the remaining Partitioning Reagent layer. The Partitioning Reagent was removed using a 200 µl tip with care to ensure that the emulsion layer was kept intact.

##### 2. Lysis of nuclei to obtain RNA and stabilize sample

Once nuclei for each sample were emulsified in the PIP tubes, we prepared the Chemical Lysis Emulsion for each PIP tube by adding 120 µl Partitioning Reagent (Fluent) to a 0.5 ml tube of Chemical Lysis Buffer 3 (#FB0003909, Fluent). The Chemical Lysis Emulsion is to be prepared as an additional emulsion that is mixed into the PIP emulsion. When the emulsions combine and are heated, the Chemical Lysis Emulsion enables lysis of the nuclei. Upon lysis of nuclei, the liberated RNA are instantly captured within each PIP droplet. To perform this step, we added 150 µl of the Chemical Lysis Emulsion to each PIP tube and gently inverted each tube to mix the 2 emulsions until cloudy (10 cycles). Each PIP tube was then placed in the 66°C digital dry bath and the heated lid of the device was set to 105°C. The Nuclei Lysis Program that is stored in the device (setting B) was then initiated. This heating regimen included 45 minutes at 66°C, 10 minutes at 25°C, and a final stage that holds the samples indefinitely at 20°C. In the current study, we removed the samples once they reached the last stage. We maintained samples at ambient temperatures (15-25 °C) and submitted them to the sequencing core within 24 hours.

#### D. Generating cDNA libraries and PIP-sequencing

In the current study, the sequencing core was the New York University Genome Technology Center (NYU-GTC). They prepared and sequenced cDNA libraries from the RNA. The procedures for cDNA library preparation were followed exactly as in the PIP-seq T2 v4.0PLUS User Guide (p. 24-43). Library preparation quality was confirmed using a Qubit Fluorometer (Invitrogen, #Q32866, ThermoFisher) to measure total cDNA yield and using a high-sensitivity TapeStation 4200 (#G2991BA, Agilent) with a HSD1000 ScreenTape (#5067-5584, Agilent) to perform the cDNA fragment analysis.

The cDNA libraries were sequenced using a NovaSeq X+ platform (Illumina) using the recommended parameters for paired-end PIP-seq with the recommended sequencing depth of 20,000 reads per nucleus (User Guide, p. 44). Flow cell quality metrics were obtained after sequencing. We verified that all samples had at least 95% of bases above a Phred algorithm quality score of 30. Quality scores above 20 are considered acceptable and a score above 30 is a common gold-standard (Ewing *et al*., 1998; Shouib *et al*., 2025). The mean quality score for each sample was between 39-40, indicating that cDNA libraries for all samples were sequenced with excellent fidelity (Ewing *et al*., 1998; Shouib *et al*., 2025).

### III. Data analysis using our pipeline

#### A. Pre-processing of PIP-sequencing data

Raw FASTQ files were obtained from the NYU-GTC for each sample. FASTQ files are a standard format for sequencing data that also includes quality scores. Next we used PIPSeeker (v3.3.0, Windows/Linux version, Fluent) to perform “cell calling”. This pre-processing step is used to assess the quality and yield of transcripts sequenced for each nucleus per sample. This allows users to compare quality of samples in terms of average metrics for all nuclei in a sample. It also enables a crucial step, in which nuclei with the highest ranked quality metrics can be selected (or “called”) to be used for all subsequent analyses. If the number of expected nuclei is unknown, 1 of 5 sensitivity tiers can be chosen to call more lower-ranked nuclei or to limit analysis to only “top-tier” nuclei. Instead, we chose to select nuclei by the expected number per sample of 5000 (via the force_cells parameter) because that is the maximum recommended nuclei input for PIP-seq T2 v4.0PLUS kits. However, it should be noted that the 5000 nuclei called for each sample are also subjected to additional quality thresholds before analysis (see below). For all PIPSeeker procedures, the relevant parameters were changed from the default to those that were most relevant to our analysis: *star_index_path* (e.g. “reference genome” used to interpret RNA sequences as segments of mouse genes) *= pipseeker-gex-reference-GRCm39-2022.04*, *threads = 4, random_seed = 0, chemistry = v4, downsample_to = none, input_reads = none, retain_barcoded_fastqs = FALSE, remove_bam = FALSE, exons_only = FALSE, force_cells = 5000, and run_barnyard = FALSE*. In contrast, default parameters were used for clustering in PIPSeeker because the clustering did not influence our downstream analyses. Notably, our approach has great flexibility of parameters, allowing users with different goals the ability to make parameters best suited for their study.

#### B. Using Seurat with SOCTS to analyze snRNA-seq datasets

##### 1. Construction of analysis pipeline

Output files obtained from PIPSeeker (6 per mouse/sample) were used as inputs to perform our snRNA-seq analyses in R (see below). These 6 inputs for R analyses consisted of 6 PIPSeeker output files that were derived from the raw (all droplets in a sample) and the filtered (5000 nuclei) datasets after processing with PIPSeeker. The PIPSeeker output file types used for the raw and filtered dataset were the matrix of read counts (“matrix”.mtx), the genes to which the reads corresponded (“features”.tsv), and the exact nuclei that contained these reads for these genes (“barcodes”.tsv). No samples were excluded from analysis.

To analyze these snRNA-seq data, we wrote a software pipeline in R (v4.4.2; R Core Team, 2024) that uses the Seurat package (v5) to quantify gene expression in specific cell types (Hao *et al*., 2024). We based our pipeline on several rigorous and widely adopted snRNA-seq tutorial recommendations. These primarily consisted of the Seurat documentation and tutorials available from both the Satija Lab website (https://satijalab.org/seurat; last accessed: 07/09/2026) and from the Harvard Chan Bioinformatics Core (HBC; https://hbctraining.github.io/main/; last accessed: 07/09/2026). Rationales and other explanations for each component of our deposited R pipeline code are available (https://osf.io/qgty7). In this section, we will explain the key parameters we used in each step of our pipeline.

##### 2. Correction for ambient RNA captured in PIPs

A common issue that emerges during nuclei isolation is that ambient RNA levels can be high. The consequences can include high variability and masking of biological effects (Caglayan *et al*., 2022). Column-based kits such as the 10x Genomics Chromium kit used in this study are often affected, but may have higher nuclei yields (Kersey *et al*., 2026). Therefore, we used the 10x Genomics Chromium kit, but corrected for the influence of ambient RNA using the R package SoupX (Young and Behjati, 2020). We set the correction to the most stringent tier (contamination fraction = 0.2) to ensure that the correction remained consistent and sufficient across samples to reduce the influence of ambient RNA.

##### 3. Thresholding to permit or exclude nuclei

After correcting for ambient RNA, we used the CreateSeuratObject() function to generate Seurat Objects from the SoupX-adjusted RNA read count datasets (1 object/mouse or 1 object/sample). In this analysis, each Seurat Object starts as a container that holds all snRNA-seq data that corresponds to one mouse (one sample), as well as all labels, metadata, and transformations performed on the data. This step is repeated for each mouse. In later steps, the 6 Seurat Objects from the 6 mice are merged into one Seurat Object to perform final analyses.

Nuclei for each sample were filtered to remove low quality nuclei using parameters recommended by the Satija Lab (https://satijalab.org/seurat/articles/pbmc3k_tutorial) and HBC materials (https://hbctraining.github.io/scRNA-seq/lessons/04_SC_quality_control.html). Nuclei were retained for analysis only if they contained over 200 transcript reads from more than 200 unique genes. To remove damaged or otherwise aberrant nuclei, another threshold was set to remove nuclei that had more than 10% of their gene expression from mitochondrial RNA (mt-RNA). A threshold of 10% mt-RNA was selected because the expected range of %mt-RNA for cells in the hippocampus is no greater than 10% in healthy mice (Osorio and Cai, 2021). In addition, the typical %mt-RNA in cells in the DG is 3-5% in baseline conditions (Osorio and Cai, 2021). Therefore, to account for variation in cells from WT and Tg2576 mice, we set the threshold greater than the upper limit of 5% in DG but did not exceed 10%. In the deposited code (https://osf.io/qgty7), this was implemented using the subset() and PercentageFeatureSet() functions to retain nuclei only if *Count_RNA > 200, nFeature_RNA > 200, and percent.mt < 10*.

Generation of one droplet containing two or more nuclei (a doublet or multiplet) is a common problem encountered in snRNA-seq data, but current methods to remove doublets have limitations. Two common approaches, using a read count threshold or an automatic detection package (such as Scrublet or DoubletFinder), were not recommended by the HBC (https://hbctraining.github.io/scRNA-seq/lessons/04_SC_quality_control.html). One primary disadvantage that was relevant to our study was that those 2 approaches often remove nuclei that are in the process of maturation or other transitional states (Petegrosso *et al*., 2020; Jiang *et al*., 2023). Therefore, there was a risk in using these approaches for our analysis because the adult DG includes GCs that undergo maturation after postnatal neurogenesis in the DG (Seki, 2011; Kempermann, 2025). Due to the risk of removing GCs, we instead chose to use the recommended approach, which was to remove cells that did not show expected combinations of marker genes. That exact procedure (i.e. for removing doublets) is performed automatically as a result of using SOCTS in our pipeline to sort nuclei (see below). A weakness of all 3 approaches is that it is difficult to detect if a doublet has been formed by two cells with identical markers for the same cell type. However, we proceeded because this weakness was unlikely to influence differential gene expression between Tg2576 and WT mice *within each cell type*. In addition, because doublet formation is a stochastic process involving random interactions of nuclei and droplets, it is highly unlikely that genotype would impact the rate at which doublets form. This would be true whether a doublet is from two nuclei of the same cell type or different cell types.

Next, the nuclei in the Seurat Object for each sample were normalized using the SCTransform() function with default parameters. SCTransform was used to normalize technical variability in the transcript counts detected in different nuclei. It does so by performing multiple steps based on a negative binomial regression of the data. This process accounts for variation between cells in sequencing depth and thereby enables valid comparisons of gene expression in pooled groups of nuclei (such as by cell type and/or genotype).

##### 4. Labeling and sorting nuclei by cell type using SOCTS

After normalizing each Seurat Object for each sample, the AddMetaData() function was used to label nuclei with the respective sample’s ID and mouse’s genotype (named “Condition”). To blind analysis, the Condition was coded as “A” or “B” instead of the genotype. Each Seurat Object was then sorted to retain nuclei of only one DG cell type (see Figure 2A) using the subset() function with logical operators (as shown in the sample of code below) to regulate the processing of data (i.e. filter). One R script was generated for each DG cell type. An example of R code used to retain only Mature GCs for analysis was:

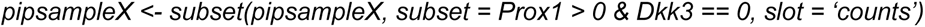

where *pipsampleX* is the Seurat Object that contains the current nuclei under analysis from one mouse and *subset()* is the name of the function performed on *pipsampleX*. The *subset =* parameter was set such that the number of transcripts detected for the positive marker gene *Prox1* would be greater than 0 and there would be exactly 0 (==) transcripts detected for the negative marker gene *Dkk3.* These marker genes are used to define Mature GCs because they express *Prox1* but do not express *Dkk3* (Cembrowski *et al*., 2016). The *slot =* parameter was set to perform the sorting (subsetting) operation using the transcript counts for each nucleus. This SOCTS sorting procedure was performed on the Seurat Objects for all 6 samples for all 9 cell types.

##### 5. Dimensional reduction and integration of dataset

It is recommended practice to perform what is called integration, in which shared statistical features of datasets from different sources, methods, or processing batches are identified (“co-registered”) and a batch correction is performed. Batch correction is necessary to enable valid comparisons when there may be batch-dependent effects (Korsunsky *et al*., 2019). Although our samples were processed in the same batch, we still used batch correction because the perfusion and sectioning of one animal might differ from another animal even if performed on the same day and processed together afterwards.

Integration and batch correction followed the gold standard using the Harmony package (Korsunsky *et al*., 2019). We first merged all samples into one Seurat Object using the merge() function. We then performed PCA using RunPCA() to generate the top 50 PCs and used them as inputs for integration. We performed Harmony integration via the IntegrateLayers() function with the following standard parameters: *assay = “SCT”, method = HarmonyIntegration, orig.reduction = “pca”, new.reduction = “harmony”,* and *normalization.method = “SCT”* (see https://satijalab.org/seurat/articles/integration_introduction).

##### 6. Clustering of nuclei using Uniform Manifold Approximation and Projection (UMAP)

For each cell type, nuclei populations were visualized via UMAP using 3 consecutive functions: FindNeighbors() with *reduction = “harmony”* and *dims = 1:15*, FindClusters() with *resolution =* minimum needed to form 2 clusters (3 for Mossy Cell UMAPs), and RunUMAP() with *dims = 1:15* and *reduction = “harmony”.* The parameters used for FindClusters() were selected to generate 2 clusters for each cell type to both visualize possible subpopulations and to demonstrate that the number of clusters generated is user-selected (see https://satijalab.org/seurat/articles/pbmc3k_tutorial). The DimPlot R package was used to generate images of UMAPs that were then added to Figures with Adobe Illustrator 2026 (v.30.5.1).

##### 7. Detecting DEGs in Tg2576 vs WT mice

Prior to performing DEG identifications, we performed pseudobulking, a common procedure. Without pseudobulking, the N used to calculate differential gene expression is the number of nuclei per condition (here, genotype). After pseudobulking, the N is the number of mice per genotype. As mentioned above, pseudobulking is performed to prevent inflation of sample sizes and thereby reduce the risk of detecting false positive DEGs (Squair *et al*., 2021; Murphy and Skene, 2022). We performed pseudobulking using the AggregateExpression() function with the parameters: *group.by = c(“Condition”, “SampleID”)* and *return.seurat = TRUE*. The active “Ident” status was set to “Condition” to compare Tg2576 to WT mice when using the FindMarkers() function with parameters: *assay = “SCT”, fc.slot = “scale.data”, ident.1 = “X.Tg2576.”, ident.2 = “X.WT.”,* and *test.use = “poisson”* (rationales for these choices are in Results, Section IID). The FindMarkers() function uses a Bonferroni correction to adjust for multiple comparisons for our DEG analyses. We used the EnhancedVolcano package to generate preliminary volcano plots within R. All volcano plots shown in the Figures, however, were recreated from the DEG list .csv output from R (1 DEG list per DG cell type in Tg2576 vs WT mice) and formatted using Prism (v10.6.1; Graphpad).

### IV. Additional analysis of outputs from pipeline

#### A. Comparisons against the standard method in Nelson et al. (2023)

We downloaded the mouse hippocampal snRNA-seq data used in Nelson *et al*. (2023) from a public repository (https://github.com/Erik-D-Nelson/ARG_HPC_snRNAseq; last accessed: 07/09/2026). We reanalyzed their datasets using our SOCTS-based pipeline to compare its performance to that achieved by Nelson *et al*. (2023). To enable a fair yet rigorous comparison, we modified our pipeline’s R code to process their datasets using identical R parameters with some exceptions described below (and see Results, Section IIC, and deposited code: https://osf.io/qgty7).

In brief, we performed our analysis of Nelson *et al*. (2023) via the following methods: the 2 ECS and 2 Sham mouse datasets were downloaded as a single SingleCellExperiment object rather than a Seurat object. SoupX was not performed because the nuclei of Nelson *et al*. (2023) were isolated via NeuN-labeled flow cytometry and SoupX was not used. We used the which() and sce.subset() functions to split the single downloaded object into 4 datasets. We converted each dataset to a Seurat object using the as.Seurat() function. Each dataset’s nuclei were then subjected to the same quality threshold filtering and labeling in our pipeline (described above). Nuclei were sorted by DG cell type using SOCTS (described in Results, Section IIC). The 4 Seurat objects were then processed as described above using the merge(), VariableFeatures(), and RunPCA() functions. Unlike Nelson *et al*. (2023), we used Harmony to perform the batch correction using the IntegrateLayers() function (as described above). Clustering and visualization steps were similar to our own pipeline, but the number of PCs used was 1:50 to match the number of PCs used in Nelson *et al*. (2023). The resolution parameters were set to either 0.1 (Mature GCs, MCs) or 0.01 (PV, SST, CCK) depending on which resolution tier achieved better separation of the most obvious clusters in each cell type. Pseudobulking and calculation of differential gene expression were performed as in our pipeline described above, with 2 exceptions. In Nelson *et al*. (2023), DEGs were identified using an uncommon method (glmQLFtest function) that performs a “Quasi-Likelihood” F-test on a log-transformed generalized linear model (GLM) with correction via Benjamini-Hochberg FDR (Benjamini and Hochberg, 1995). We instead used one of the most common methods, DESeq2 (Love *et al*., 2014), which is not substantially different than the method used by Nelson *et al*. (2023) as both use a log-transformed GLM. However, our method corrected for multiple comparisons using a more conservative method than Nelson *et al*. (2023), the Bonferroni correction.

#### B. Identifying the overlapping genes between various DEG lists

In this study, we identify shared or unique genes between various lists from different sources (cell types, species, study methodologies). We performed these overlap analyses in R using the VennDiagram package with the function *calculate.overlap()* to generate respective lists in a .csv file of intersecting or unique DEGs for respective comparison. When results were displayed as Euler plots (as in Figure 3), the online application for the EulerR package (currently named “Eunoia”; https://eunoia.bz/; last accessed: 07/09/2026) was used to generate the Euler plots. When results were displayed as upset plots (as in Figure 6), the online application for the UpSetR package (https://gehlenborglab.shinyapps.io/upsetr/; last accessed: 07/09/2026) was used to generate the upset plots.

### C. Gene Ontology of biological functions related to DEGs

In Python (v3.11.11: https://python.org; last accessed: 07/09/2026), the package Ontomancer was used to characterize the functions that may be impacted by alterations in gene expression. It uses gene set enrichment scoring akin to methods like Integrated Pathway Analysis and performs ontology analysis using multiple ontology databases that can be obtained from GSEA (e.g. GO Biological Function, KEGG, REACTOME; Subramanian *et al*., 2005). Ontomancer is currently used by the laboratory of Dr. Avtar Roopra to perform ontology analyses and has already been used in peer-reviewed studies (Hoffman *et al*., 2025). A publication describing Ontomancer is currently in preparation by the Roopra Lab and code will be made available via GitHub when released. For this reason, Dr. Roopra kindly performed these specific analyses (in Figure 7) by passing our DEG lists into Ontomancer and obtaining the outputs corresponding to ontology analysis by the REACTOME database. Fisher Exact Tests (corrected with Benjamini-Hochberg FDR) were used to calculate the overlap between the gene set that corresponds to a given biological function and the list of DEGs (Benjamini and Hochberg, 1995).

#### D. Analysis to identify putative transcriptional regulators of DEGs

In Python (v3.11.11; https://python.org), we used the package MAGIC (Mining Algorithm For Genetic Controllers; https://github.com/asroopra/MAGIC; last accessed: 07/09/2026) to identify possible upstream TFs that could be driving alterations in expression of target genes in our DEG lists. MAGIC was used to rank TFs with the highest proportions of their respective target genes that overlap with genes on a given DEG list. It uses target gene lists from the human ENCODE TF database of chromatin immunoprecipitation (ChIP) experiments (Luo *et al*., 2020). It performs these calculations as described in the original study that introduced MAGIC (Roopra, 2020). Nonparametric Kolmogorov-Smirnov tests corrected with Benjamini-Hochberg FDR were used to calculate significant statistical enrichment of DEG lists with TF target gene lists. This method also accounts for the number of all genes expressed (the “background”, DEG or not) in each analysis in comparison to numbers of DEGs to reduce the chances of false positives. A MAGIC Score is also calculated alongside the significance of enrichment that allows TFs to be ranked by corrected p-values with an adjustment that selects for TFs with robust binding signatures in the ENCODE database (Figure 7A-B). For each TF, the strengths of binding at gene loci were derived from the ChIP experiments used to construct the ENCODE database (Luo *et al*., 2020).

### V. Additional details of statistical methods

For all analyses, the thresholds of statistical significance for the adjusted-p or FDR values were: α = *0.05, **0.01, or ***0.001. A two-tailed distribution was used for all statistical tests that required selection of tails. Unless otherwise stated, all analyses were performed and graphics generated using GraphPad Prism v10.6.1. In bar graphs, data are either shown as mean ± SEM when single values indicate the number of DEGs (Figures 3D and 6B-J), or an odds ratio (Figure 7C-D). As described above, SCTransform was used for normalization of snRNA-seq transcript count datasets. Calculations for detection of DEGs are described above (Methods, Section IIIB-7 and Section IVB). Normality was assessed for datasets in Figures 2, 3, and S3 using the Shapiro-Wilk and Kolmogorov-Smirnov tests. In addition, Spearman’s test of heteroscedasticity was used to ensure that each dataset was homogeneous in variances so that parametric statistics could be performed. Data in Figure S3 were normalized as percentages in order to perform parametric statistics on the data shown in Figure 2C. In Figures 2C and 3C, a two-way ANOVA was used to perform a comparison of means for more than 2 groups and was corrected for multiple comparisons using the Bonferroni method to reduce risk of Type I errors. Except for Figure 7, all statistical tests involving post-hoc multiple comparisons were corrected using the Bonferroni method, as it is generally more conservative. In Figure 7A-B, outputs from MAGIC analyses are presented. Because the data were non-parametric, MAGIC performed Kolmogorov-Smirnov tests corrected with Benjamini-Hochberg FDR (Benjamini and Hochberg, 1995) on each DEG list to calculate which TFs showed significant enrichment (e.g. high overlap of a TF’s target genes with DEGs). A MAGIC Score was calculated in which TFs were ranked by corrected p-values with an adjustment that selects for TFs with the most robust binding signatures. For Ontomancer analyses in Figure 7C-D, data were parametric and therefore Ontomancer used a Fisher’s Exact Test that it automatically corrected using Benjamini-Hochberg FDR. In addition, a Fisher’s Exact Test was used for comparisons of the observed vs expected proportions of DEGs in Figures 3D-F or nuclei with correct cell type labels in Figure S5. We selected the Fisher’s Exact Test instead of a Chi-square test because the risk of Type II errors is robustly increased when the Chi-square test is used to compare small (< 5-10) numbers, which were present in some of our comparisons. In contrast, the Fisher’s Exact Test can compare large or small numbers without profound loss of statistical power (Lewis and Burke, 1949).

## Supporting information

DataS1 DEG lists for all panels in Fig 4

DataS2 Gene list sets related to Fig 6

STable1 SOCTS table and rationales

STable2 Outputs from MAGIC and Ontomancer in Fig 7

## SUPPLEMENTAL FIGURE LEGENDS

**Figure S1.**
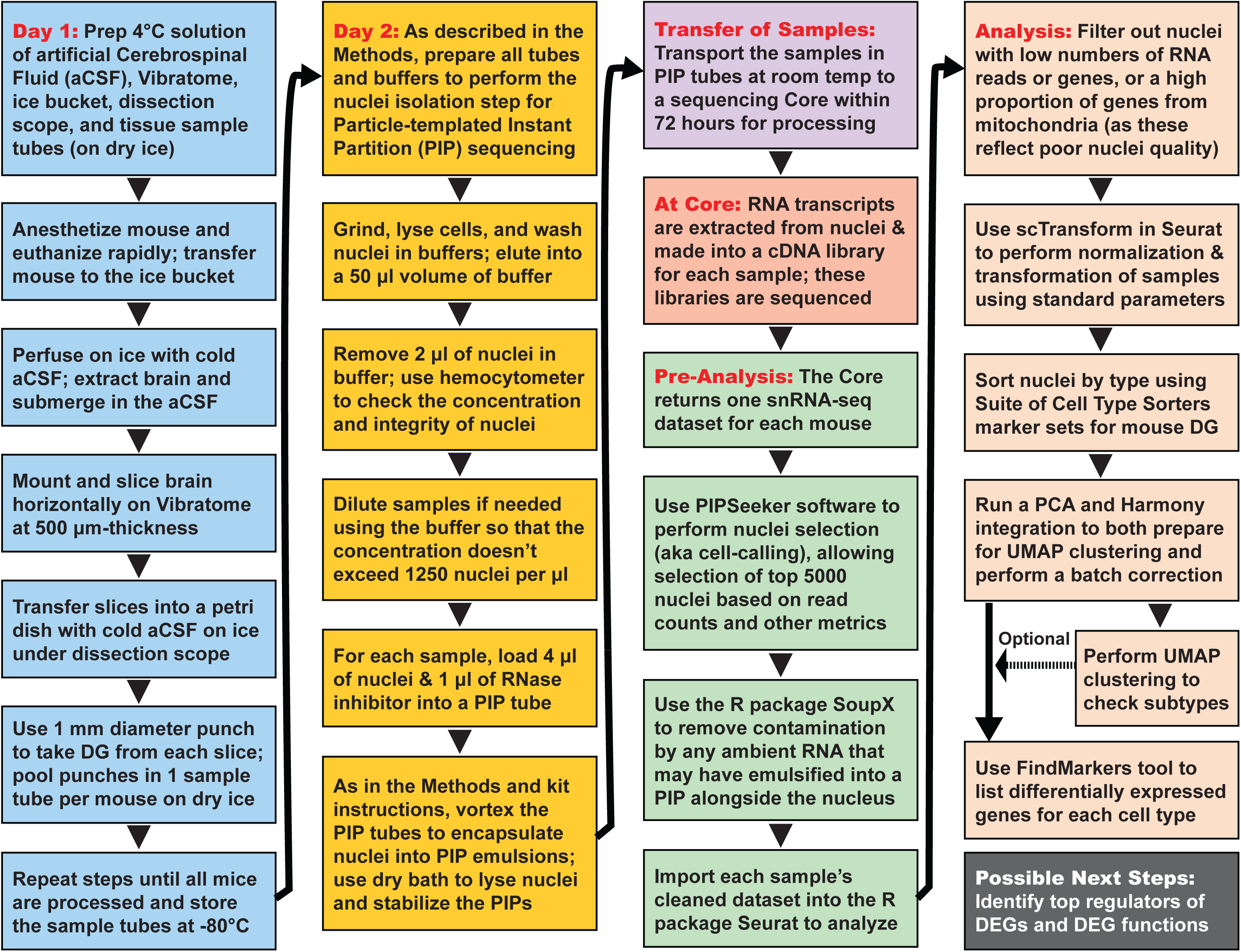
Overview of snRNA-seq and analysis pipeline. Refer to Methods for additional description.

**Figure S2.**
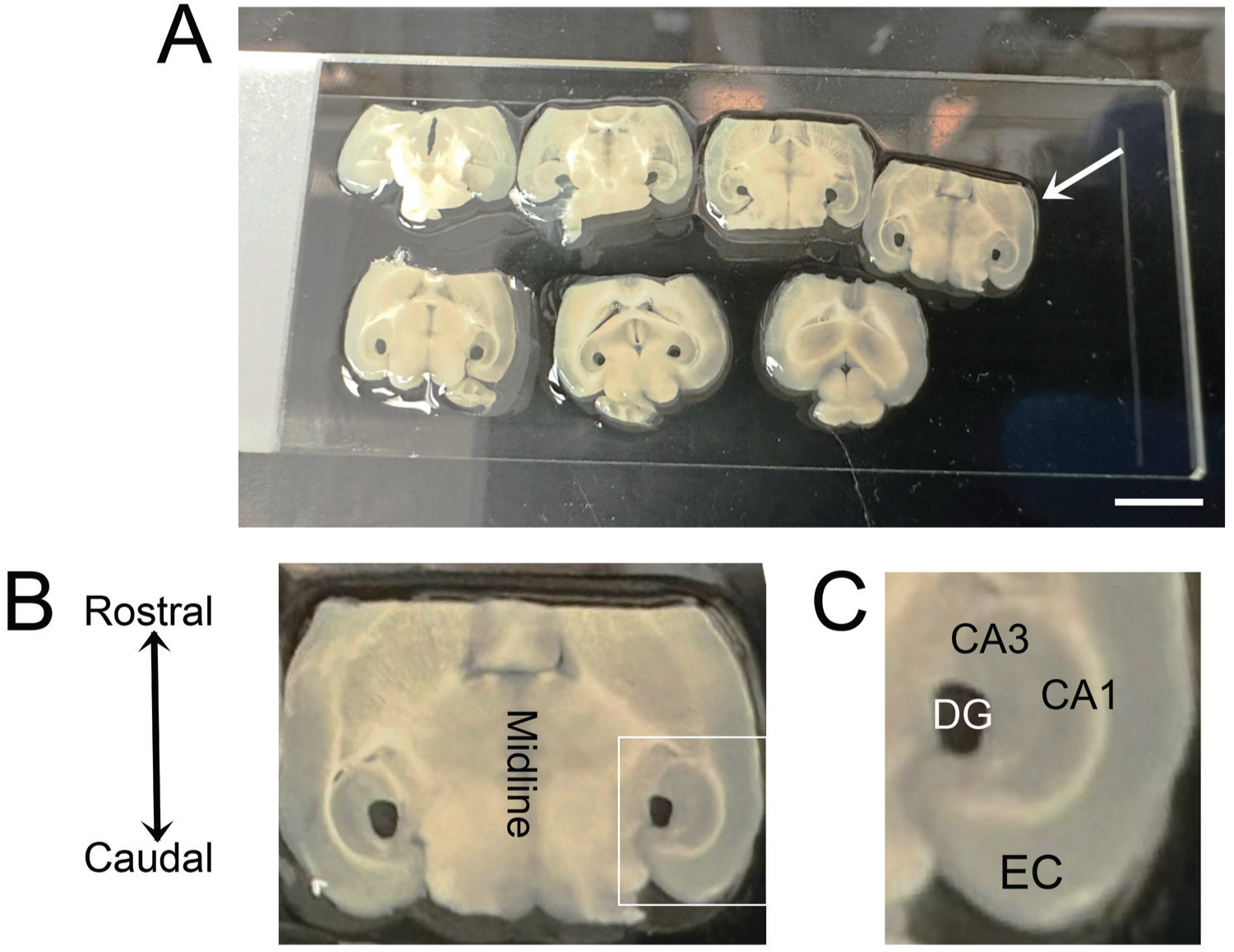
Anatomical location of tissue harvested from DG. **(A).** An example of slices from 1 mouse, arranged in order from most dorsal to most ventral, with the biopsy punches shown. The white arrow indicates the slice that is enlarged in (B-C). Scale bar = 5 mm. **(B).** The midline and the rostral-caudal axis are shown. **(C).** The white box in (B) is enlarged. DG, dentate gyrus; entorhinal cortex, EC; area CA1, CA1; area CA3, CA3.

**Figure S3.**
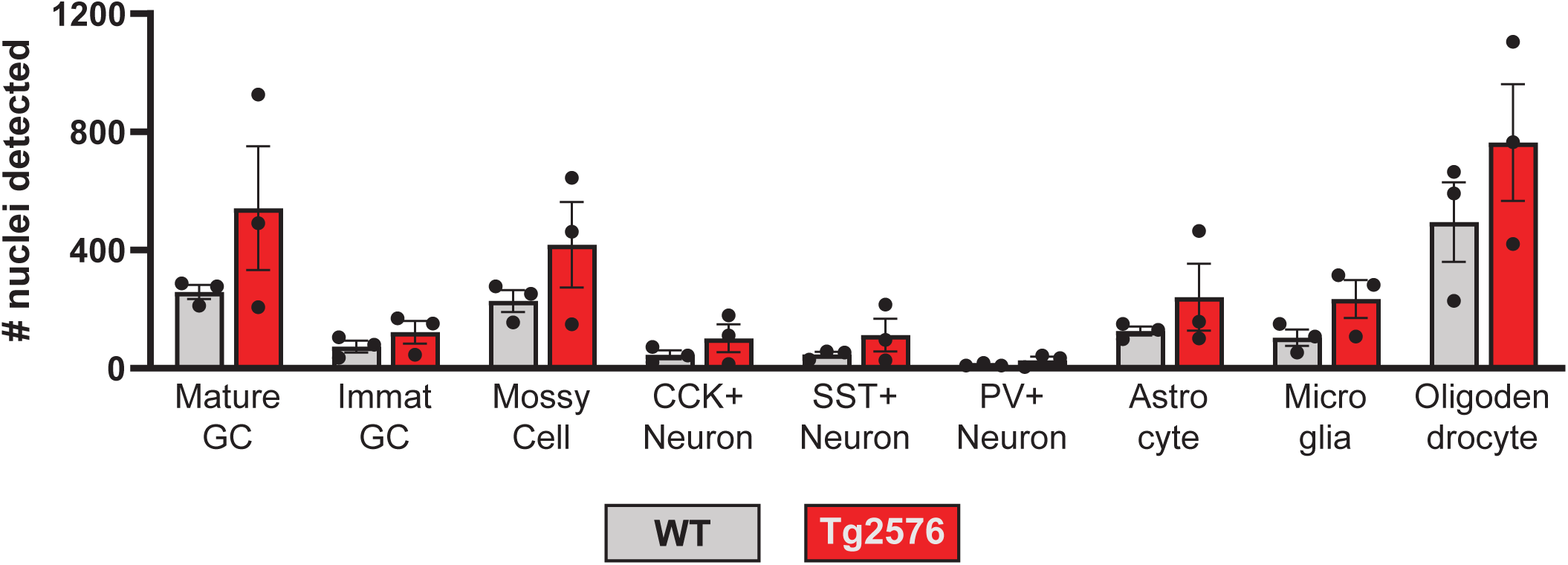
Numbers of nuclei detected for each DG cell type in Tg2576 and WT mice. In Figure 2C, proportions of nuclei per cell type are presented. We show here the raw numbers of nuclei detected per cell type in Tg2576 or WT mice. Note that statistical comparisons are in Figure 2C rather than here because the raw data showed significant departure from normality (Shapiro-Wilk: W = 0.8890, p = 0.0001; Kolmogorov-Smirnov: distance = 0.1866, p < 0.0001).

**Figure S4.**
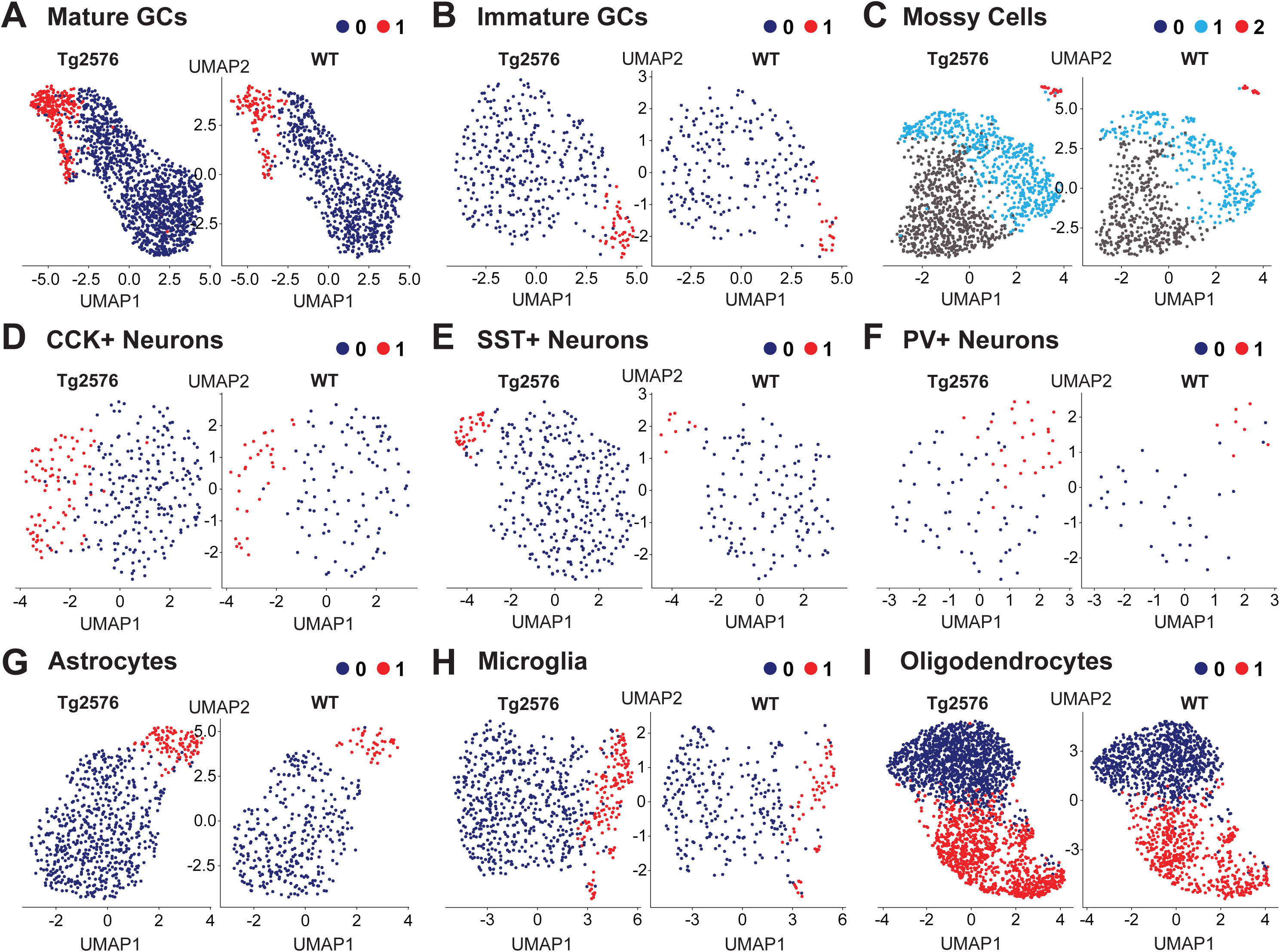
Clustering of nuclei for individual DG cell types in Tg2576 and WT mice shows possible subtypes. **(A-I).** Standard UMAP clustering was performed to visualize the nuclei populations obtained for each DG cell type in Tg2576 and WT mice. As detailed in the Methods, UMAP analyses were performed on principal components obtained from PCA of nuclei in each cell type to generate 2 clusters for each cell type. The DimPlot package in R was used to generate images of UMAP clusters. One plot was obtained for each genotype for each DG cell type, with Tg2576 UMAPs on the left and WT UMAPs on the right of each panel. Cluster names were simply “0”, “1”, and “2”, assigned by default, and are distinguished by colors. Cluster names were left unchanged from the numerical names because they were intended to not imply cell type, reducing bias. Note multiple clusters are evident, suggesting subtypes. The subtypes could reflect, for example, differences in dorsal vs. ventral MCs since they have been demonstrated (Cembrowski *et al*., 2016; Botterill *et al*., 2021; Fredes *et al*., 2021; Houser *et al*., 2021; Jeong *et al*., 2026).

**Figure S5.**
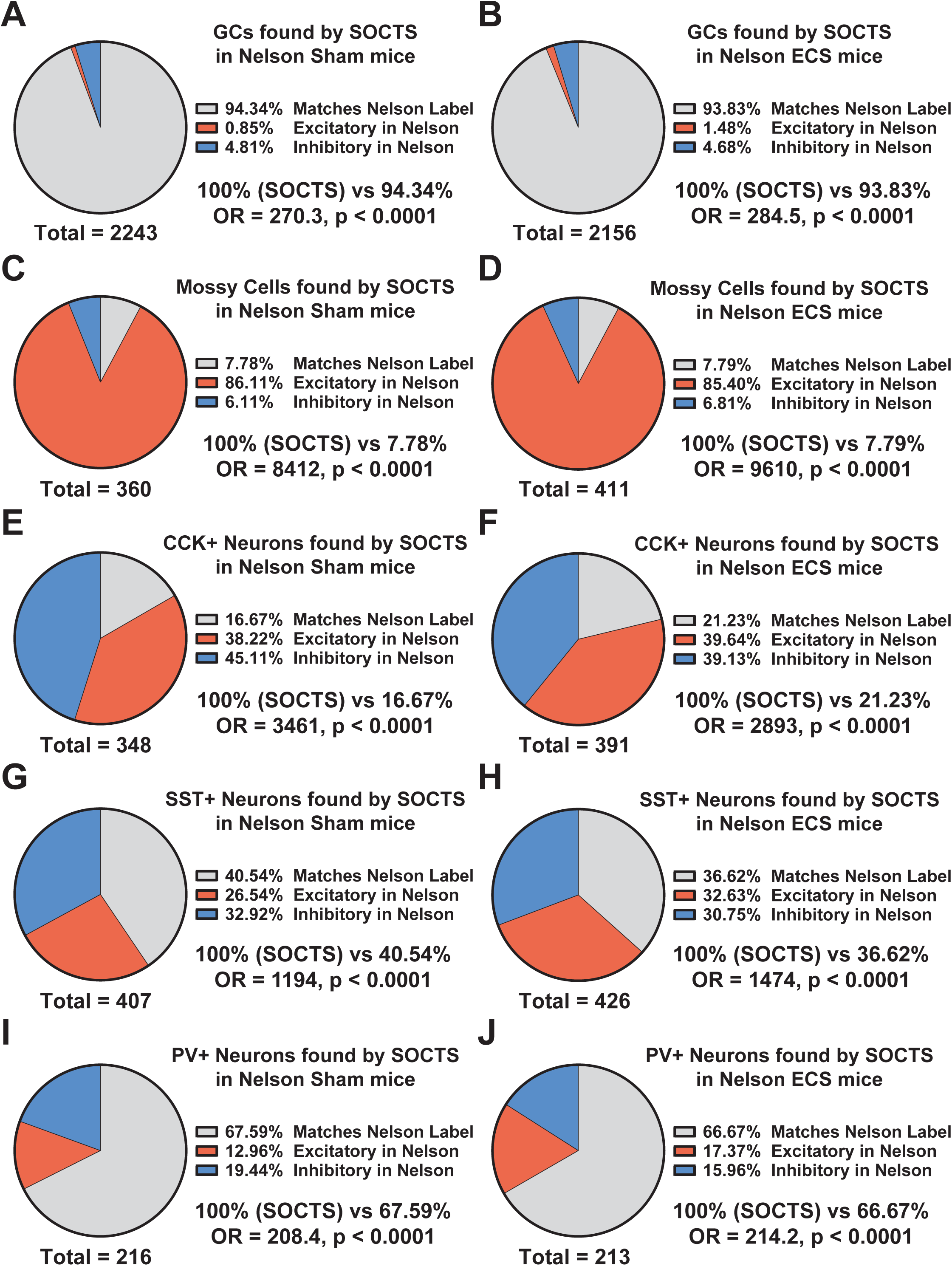
Standard methods can result in inconsistencies in identification of cell types. In Figure 3C, we show the number of nuclei that were detected for each neuronal cell type in a standard approach (i.e. Nelson *et al*. 2023) compared to when we used SOCTS. In Figure S5, we analyzed the proportion of cell type labels in Nelson *et al*. (2023) that were consistent with the cell type labels applied by SOCTS. The marker gene sets used by SOCTS were in alignment with validated marker genes that are used to confirm cell types in the DG (Table S1) and therefore are represented by 100% in A-J. **(A-J).** We tested the specificity of the Standard approach by comparing if the original cell type labels aligned with the labels applied by SOCTS. We performed this analysis on the Sham and ECS groups (n = 2 mice/group) separately to ensure cell labeling agreement did not depend on treatment condition. The nuclei labeled by SOCTS that do not match the cluster labels in Nelson *et al*. (2023) are displayed as “Excitatory in Nelson” (*red*) or “Inhibitory in Nelson” (*blue*). The comparison illustrates that there is disagreement in the approaches, suggesting inconsistencies in the output of the Standard approach. Significance was calculated using Fisher Exact Tests to compare the proportions of cell type labels that were applied by SOCTS vs the Standard approach that were consistent or inconsistent with the marker genes expressed by each nucleus. In all 5 cell types, the proportions of nuclei that matched the expected marker genes for the cell type were significantly greater when using SOCTS vs the Standard approach (***p < 0.0001). For each method, the proportions of nuclei that were assigned labels consistent with their marker genes are reported with the odds ratio (of the difference between methods) and p-value. The “total =” shown below each diagram refers to the total number of nuclei that were identified by SOCTS for each neuronal cell type after reanalysis of the data from Nelson *et al*. (2023). Abbreviations: Excit, excitatory, Inhib, inhibitory; SOCTS, suite of cell type sorters; UMAP, uniform manifold approximation and projection; ECS, electroconvulsive shock.

**Figure S6.**
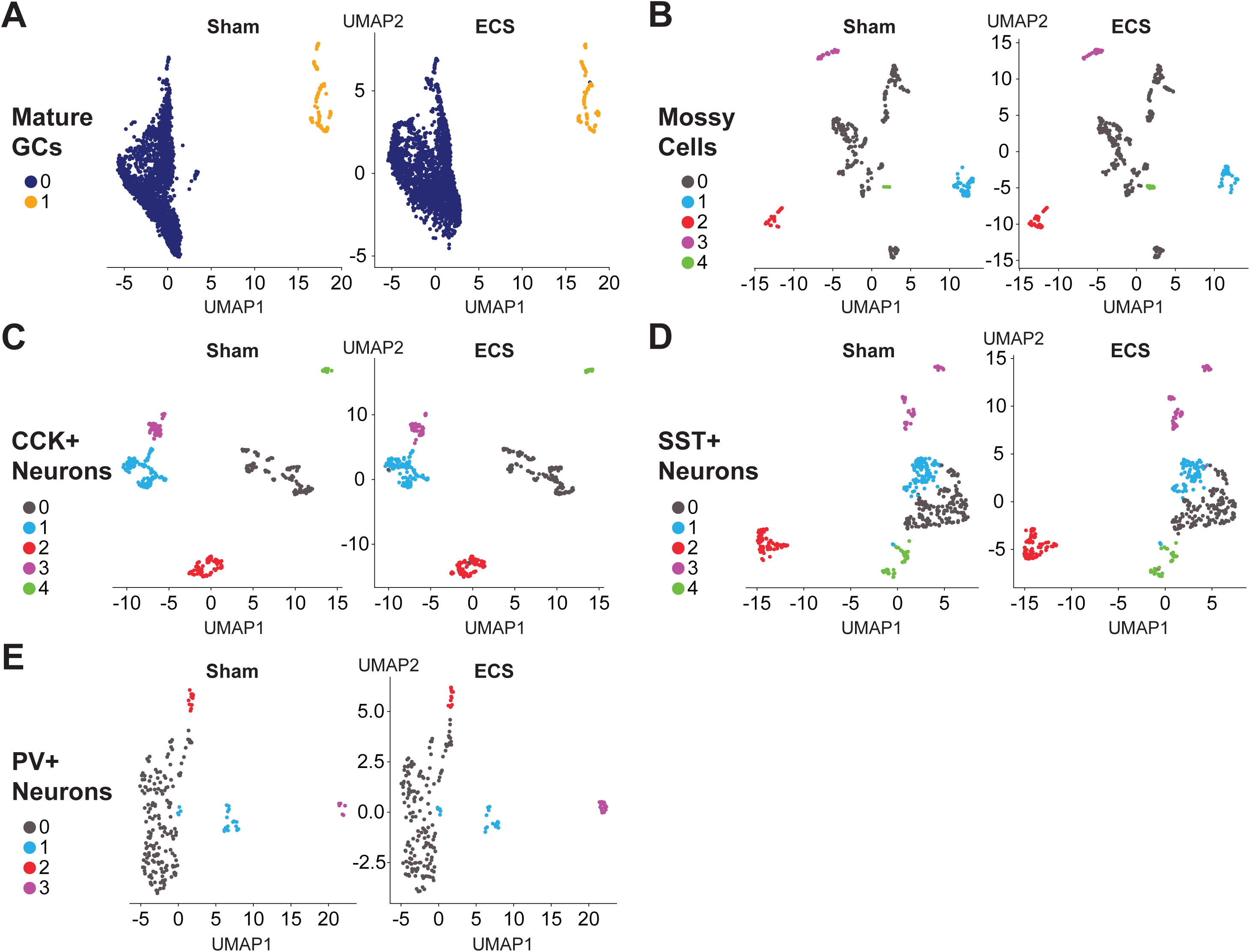
Clustering of nuclei in Nelson *et al*. (2023) can be performed after cell type identification by SOCTS. **(A-E).** We demonstrate that clustering can be performed on published snRNA-seq datasets after they are analyzed using our SOCTS-based pipeline. UMAP results are displayed in a similar manner as UMAP results in Figure S4 (see Figure S4 legend for details on UMAP plotting; refer to Methods for parameters and rationale of selections).

## CRediT AUTHORSHIP CONTRIBUTION STATEMENT

**Gabriel S. Stephens:** Conceptualization, Data curation, Formal analysis, Investigation, Methodology, Project administration, Resources, Software, Validation, Visualization, Writing – original draft, Writing – review and editing. **David Alcantara-Gonzalez:** Investigation, Methodology, Resources, Validation, Writing – review and editing. **Helen E. Scharfman:** Conceptualization, Funding acquisition, Methodology, Project administration, Resources, Supervision, Writing – review and editing.

## USE OF AIMAL MODELS

All experimental procedures were performed in compliance with NIH guidelines and approved by the Institutional Animal Care and Use Committee at the Nathan S. Kline Institute for Psychiatric Research. This study conforms with the United States Public Health Service’s Policy on Humane Care and Use of Laboratory Animals, and also with the ARRIVE guidelines for ethical reporting of animal research.

## DECLARATION OF GENERATIVE AI USE

Artificial intelligence (AI) was not used to perform any part of this project. Generative AI was also not used at any point in the generation or editing of this manuscript.

## ACKNOWLEDGEMENTS

The authors would like to thank the New York University Genome Technology Center for performing the pre-processing of RNA samples and the sequencing reported in this study, as well as their advice and guidance throughout the project. In addition, we are grateful to Dr. Avtar S. Roopra (University of Wisconsin-Madison) for his guidance and for help with the Ontomancer analyses in Figure 7C-D, as well as Jose E.C. Espina (University of Wisconsin-Madison) for help in the early stages of this study. We thank scientists at Fluent for their assistance in developing buffers compatible with our approach. We would also like to thank Dr. Stephen S. Ginsberg (Nathan S. Kline Institute and NYU) and members of the Scharfman Lab. Furthermore, we wish to recognize the animal facility staff at the Nathan S. Kline Institute for their excellent work in caring for the animals used in this project.

## APPENDIX A. SUPPLEMENTARY MATERIAL

[Journal will link to the content that contains Tables S1-S2 and Data S1-S2.]

## DATA AVAILABILITY

All underlying data and code used in the preparation of this study are freely available through the Open Science Framework (OSF) at the Center for Open Science by following this link: https://osf.io/qgty7

